# Realistic modeling of ephaptic fields in the human brain

**DOI:** 10.1101/688101

**Authors:** Giulio Ruffini, Ricardo Salvador, Ehsan Tadayon, Roser Sanchez-Todo, Alvaro Pascual-Leone, Emiliano Santarnecchi

## Abstract

Several decades of research suggest that weak electric fields may influence neural processing, including those induced by neuronal activity and recently proposed as substrate for a potential new cellular communication system, i.e., ephaptic transmission. Here we aim to map ephaptic activity in the human brain and explore its trajectory during aging by characterizing the macroscopic electric field generated by cortical dipoles using realistic finite element modeling. We find that modeled endogenous field magnitudes are comparable to those in measurements of weak but functionally relevant endogenous fields and to those generated by noninvasive transcranial brain stimulation, therefore possibly able to modulate neuronal activity. Then, to evaluate the role of self-generated ephaptic fields in the human cortex, we adapt an interaction approximation that considers the relative orientation of neuron and field to derive the membrane potential perturbation in pyramidal cells. Building on this, we define a simplified metric (EMOD1) that weights dipole coupling as a function of distance and relative orientation between emitter and receiver and evaluate it in a sample of 401 realistic human brain models from subjects aged 16-83. Results reveal that ephaptic modulation follows gyrification patterns in the human brain, and significantly decreases with age, with higher involvement of sensorimotor regions and medial brain structures. By providing the means for fast and direct interaction between neurons, ephaptic modulation likely contributes to the complexity of human function for cognition and behavior, and its modification across the lifespan and in response to pathology.

## 1 INTRODUCTION

Jefferys (*1*) defined population electric field effects as those “in which the synchronous activity of populations of neurons causes large electric fields that can affect the excitability of suitably oriented, but not closely neighboring, neurons”. The literature refers to these, loosely, as “ephaptic interactions”. Traveling at the speed of electromagnetic radiation, self-generated ephaptic fields provide the means for fast and direct interaction between neurons, enabling new mechanisms for communication and computation that remain incompletely understood. Although much faster than chemical synaptic transmission and with a longer range than electrical synaptic communication in gap junctions, electromagnetic waves travel slower in biological media than in vacuum. Table S1 in Supplementary Materials summarizes the relevant electromagnetic properties of tissues in the brain, including propagation velocity.

Work in the last decades has shown that neuronal circuits are surprisingly sensitive to weak endogenous or exogenous low frequency (0–100 Hz) electric fields (> 0.1 V/m). For example, Frohlich et al (*2*) showed that exogenous direct current (DC) and low frequency alternating current (AC) electric fields modulate neocortical network activity in slices with a threshold of 0.5 V/m. They also found effects from the application of exogenous fields mimicking endogenous fields recorded from the slices. More recent research has further established the role of ephaptic interactions and the sensitivity of neuronal populations to weak fields both in-vitro and in-silico. In particular, it demonstrates that ephaptic fields are capable of mediating the propagation of self-regenerating slow (~0.1 m/s) neural waves (*3, 4*) and that externally applied extracellular electric fields with amplitudes in the range of endogenous fields are sufficient to modulate or block the propagation of this activity both in vitro and in silico models (*5*). Field amplitudes in the range of 0.1–5 V/m have also been shown to produce physiological effects in primates using transcranial electrical current stimulation (see, e.g., (*6*) for recent results in nonhuman primates). Table S2 in Supplementary Materials provides an overview spanning six decades of in-vivo and in-vitro research on the physiological impact of weak, low frequency (< 100 Hz) electric fields—both exogenous and endogenous.

Here we focus on endogenous fields that may contribute to short-range communication at or above millimeter scales, that is, not ultra-local ephaptic effects coupling adjacent neurons. The generation of fields capable of effectively bridging such distances necessitates the synchronized activity of neuronal populations (*7, 8*) radiating from cortical patches, which occurs at frequencies below about 100 Hz (the “EEG regime”) and with spatial correlation scales in the order of a centimeter. We will call these slow, macroscopic ephaptic fields SEFs for short. As SEFs appear to be of physiological relevance (v. Table S2) and not simply an epiphenomenon, understanding how and where they play a functional role may be necessary for the development of realistic models of neural dynamics and function. Additional motivation for this study derives from seeking a theory for the effects of the weak exogenous electric fields—such as the ones generated by transcranial electrical current stimulation (tCS or tES, as it is sometimes known). At the frequencies of interest here (<100 Hz), both endogenous and exogenous tCS fields are characterized by relatively large spatial correlation scales (of the order of centimeter or more) and low magnitudes (> 0.1 V/m). Gaining a better understanding of ephaptic effects may shed some light on how tCS modulates neural dynamics and, eventually, how to optimize it.

First, we use modern biophysical modeling tools to characterize macroscopic ephaptic fields (i.e., spatially averaged at linear scales >0.1 mm, *v*. (*9*), section 4.3) using realistic head modeling. In the Methods section, we describe how we model the electric fields from EEG generating cortical populations at experimentally observed densities and patch sizes and compare them with those described in available experimental work. We analyze this first in an idealized analytical model, then in a simple 3D model, and, finally, in a realistic brain model derived from an individual MRI.

Based on this, we propose an **ephaptic modulation index** that can be computed on individual from realistic brain models (EMOD) to characterize ephaptic coupling in an individual’s brain and a derive a first simplified version for computational convenience (EMOD1). Although existing metrics such as gyrification, cortical thickness or surface area capture some geometric aspects relevant to ephaptic coupling, we take a more physics-grounded approach. We build on existing models for the interaction of weak electric fields and neurons as used in the field of transcranial current electrical stimulation (the “lambda-E” model (*10*)). Considering the cytoarchitecture of the cortex placing pyramidal cells oriented perpendicular to the cortical surface, the lambda-E model indicates that the quantity of relevance to study electric field effects is the normal or orthogonal component of the field to the cortex (*E_n_*).

Finally, we analyze how EMOD1 changes across the lifespan by characterizing it from individual structural MRIs of a large sample of 401 heathy individuals aged 16–83. EMOD1 and structural morphologies such as cortical thickness, surface area and gyrification, were correlated with age, providing a map of brain regions whose potential for ephaptic transmission is significantly affected by aging. Such findings suggest that ephaptic modulation might have relevance for cognitive processing and for the manifestation of pathological conditions involving brain morphometric changes as well alterations of oscillatory patterns (e.g., schizophrenia (*11*), depression (*12*), Alzheimer’s Disease (*13*) or Parkinson’s (*14*)).

## RESULTS

### 2.1 Ephaptic map from cortical patch sources in simplified 3D model

Median sulcal width in human brains across the age span can vary between 0.5 and 5 mm (*15*). Using this as a reference, we first studied the characteristics of endogenous fields in a 3D toy model of a sulcus in the cortex. The electric field distribution in the simplified 3D models for a sulcus width of 1 mm is shown in Figure 2 for the multiple dipole model (middle row, figures c-d) and the single dipole model (bottom row, figures e-f). Dipole strength in the multiple dipole model was set to 0.39 and 0.78 nAm, which results in a dipole strength density per unit area of 0.5 and 1.0 nAm/mm^2^ in the modeled 60 mm^2^ cortical patch. The dipole strength in the single source model was set to the same value, which results in a physiologically realistic (*16, 17*) local density of 0.5 and 1.0 nAm/mm^2^ in the equivalent area associated to this dipole (60 and 77 mm^2^). As can be seen in the figure, in the models with the higher dipole density (1.0 nAm/mm^2^), an electric field >0.1 V/m can be observed in the wall opposite to the one where the sources are located. This is observed in both the multiple and single source models, although, as expected, the area in which the electric field is greater than 0.1 V/m is higher in the former than in the latter (the electric field from multiple-source patches decays much slower than the single dipole source case (*7*), p. 37). This effect was only observed in the model with sulcus width of 1 mm. Increasing the sulcus width led to lower electric field values on the opposite sulcal wall. Figure 1 in Supplementary Materials displays the decay of the normal component of the electric field and the electrostatic potential with distance. The decays of *V* and *E_n_* are well fit by a power function with exponents of −0.66, −0.88 and −2.11, −3.02, respectively, for the multiple source and single source models.

**Figure 1:**
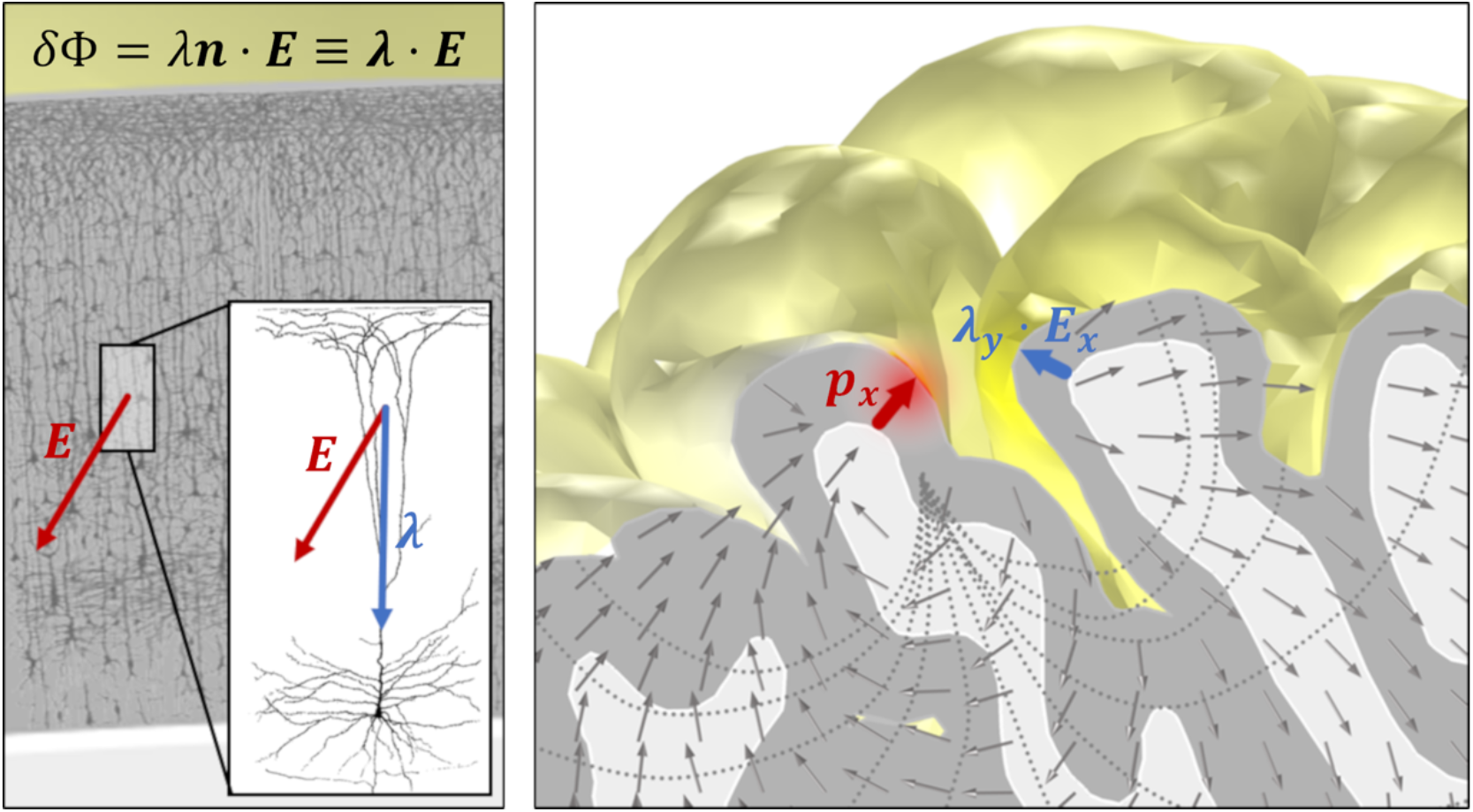
First order model for interaction of electric fields with elongated neurons. On the left, pyramidal neuron population from the human cortex (edited from “Comparative study of the sensory areas of the human cortex” by Santiago Ramon y Cajal, published in 1899, Wikipedia Public Domain). On the right, realistic model of the electric field generated by a current a dipole located at *x* in the cortex. The orientation of the generating dipole or neuron population and the sensing population (at point *y*) both play a role.

### 2.2 Ephaptic map from cortical patch sources in realistic head model

Next, we analyzed the electric fields in a realistic head model. For each one of 112 single dipole models, we calculated the decay of *E_n_* with Euclidean distance to the source. For all models, the decay was well fit by a power function, with an exponential of −3.2±0.8 (*R*^2^ of fit was 0.76±0.12). Comparing the decay of the normal component of the electric field with distance in the cortical surface, we see that it is approximately monotonic for the Euclidean distance, as expected, but not for the geodesic distance (see Figure S1 in Supplementary Materials, bottom). This behavior is expected and a result of surface folding.

For the multiple dipole source patch model, different configurations were tested using 133 dipole node sources, with individual dipole strengths adjusted so that the dipole strength area density was of 0.5 nAm/mm^2^ or 1.0 nAm/mm^2^. This resulted in individual dipole strengths at each node between 1.9 and 4.0 nAm. We also calculated single node dipole versions of these models, with strengths of 2.1 and 4.2 nAm, which correspond to the same density values in the equivalent (mesh triangle) patch size covered by that dipole. For these source strengths, it is possible to achieve an electric field magnitude of at least 0.1 V/m on the opposite sulcus wall (see Figure 3). This effect is local and dependent on the distance between source and sulcus wall. Using single source models positioned in the narrow part of the sulcus (Figure 3 bottom. b/e) and in the wide part of the sulcus (Figure 3 bottom, c/f) we found that only the former induced a 0.1 V/m electric field on the opposite sulcus wall. These results mimic closely those observed in the simplified volume conductor model discussed previously, since for the chosen study area sulcus separation in the realistic model was 1.4–5.5 mm in the dipole patch region (see Figure S2 in Supplementary Materials). For reference, sulcus width in the human cortex can be less than 1 mm (*15*). Table 1 summarizes the maxima of the electrostatic potential (at scalp level) and the electric field in the GM for all the realistic head the models presented here. See also Figure S10 for the scalp potential map associated to the chosen dipole patch.

**Table 1:**
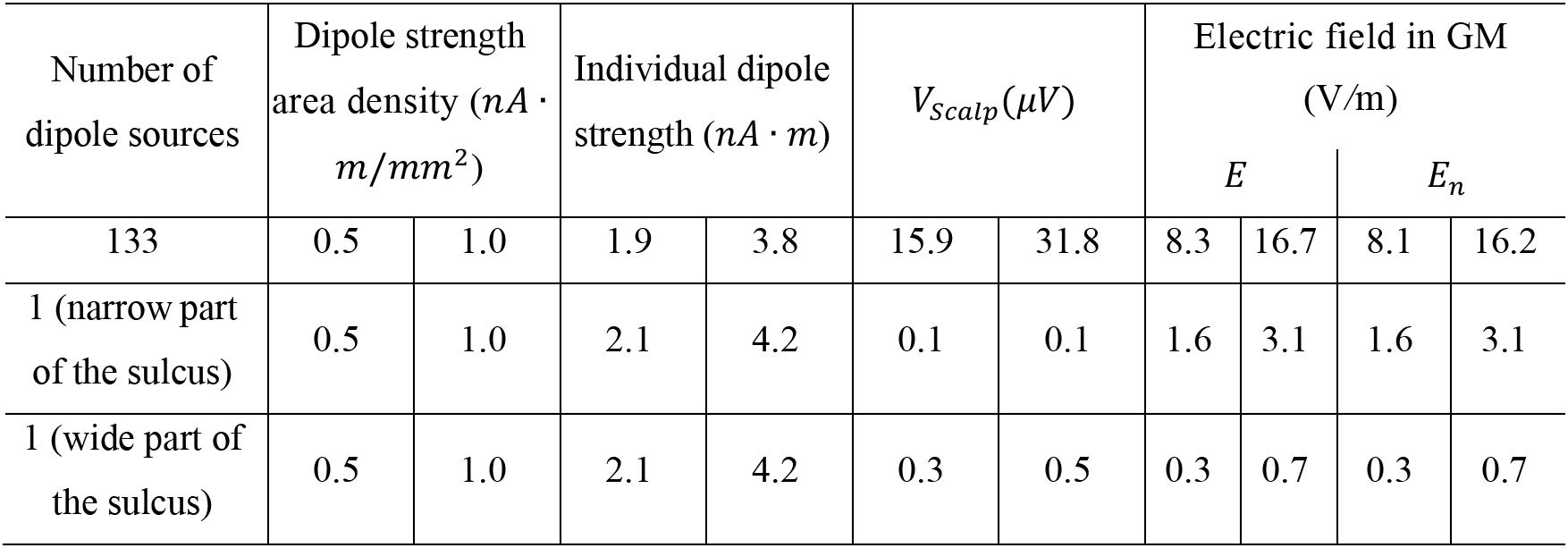
Summary of the maximum values of the scalp electrostatic potential (*V*) and GM electric field (magnitude, *E*, and normal component, *E_n_*) induced in all the source distributions used in the realistic head model. For each quantity, two dipole densities are considered: 0.5 and 1.0 nAm/mm^2^.

Finally, as a check of the realistic model, we investigated the voltage distribution at the scalp induced by a single source dipole on the chosen cortical area with a strength of 100 nAm, which is what reciprocity considerations predict would be required to achieve ~10 μV at scalp level (see Methods). The dipole was aligned to the electric field induced in that node by a montage with CP2 as the anode (1 mA) and T10 as the cathode (−1 mA). The potential difference between electrodes CP2 and T10 was of 13 μV (within the expected bounds of the approximation).

### 2.3 Ephaptic modulation in the human brain

In order to provide a template map for the distribution of ephaptic modulation in the human brain, as well as for its aging-related trajectory, 401 structural MRIs of healthy participants aged 16-83 yrs. were processed using Freesurfer software, obtaining vertex-wise cortical thickness, surface area and gyrification LGI maps for each brain. Pial surfaces obtained via Freesurfer were then used to calculate ephaptic modulation using the EMOD1 coefficient (Equation 8 with *l*_0_ = 5*mm*). A first average ephaptic map was obtained by averaging the resulting 401 EMOD1 maps (Figure 4A, Figure S4). As expected, following cortical gyrification patterns, the topography of EMOD1 displayed higher values along the sulci walls as well as medial regions such as the precuneus, and anterior cingulate cortex (see figures for statistical results).

In order to understand the relationship between ephaptic and other cortical morphologies (i.e., cortical thickness, surface area, gyrification), vertex-wise correlation was performed between EMOD1 and each morphological metric (Figure 4B). EMOD1 displayed significant but spatially different correlations with all the three morphologies, suggesting the magnitude of ephaptic modulation as potentially resulting from different cortical, non-exclusive structural patterns. EMOD1 also displayed a positive correlation with gyrification and surface area, and a negative correlation with cortical thickness following sulcal patterns (Figure 4B).

### 2.4 Changes in Ephaptic Modulation with Aging

Vertex-wise correlation between EMOD1 and age produces a bilateral pattern involving primarily sensorimotor regions, insular cortex and anterior cingulate cortex (Figure 5A). The same correlation was performed for thickness, gyrification and surface area. Globally, all metrics show a tendency to decrease with age. The decrease is very well approximated by a linear function for the EMOD1, average LGI and average thickness metrics, with *R*^2^ values of the linear fits of 0.34, 0.36 and 0.44, respectively. All of these fits are statistically significant, with p-values of 3.7 × 10^−38^, 6.5 × 10^−41^ and 9.6 × 10^−53^, respectively. For the total cortical area, the fit is worse (*R*^2^ of 0.19) but still statistically significant (p-value of 4.1 × 10^−20^). Pearson-correlation coefficients between EMOD1 and average LGI/thickness are also relatively high (0.52 and 0.43, respectively).

## 3 DISCUSSION

Understanding the functional role of ephaptic mechanisms can, among others, shed new light on the mechanisms underlying neuronal oscillations or help drive the design of better brain stimulation solutions. Research can be guided by focusing on the main features of ephaptic interactions: very fast, bidirectional, propagation of information (see Table S1) between cortical sites, influencing both local and synaptically distant regions as long as they are near in (3D) space, and in a direction dictated by the state and orientation of the emitting and receiving populations (i.e., with effects that can be both excitatory and inhibitory). For example, ephaptic interaction may play an important role in cortical recurrent computation, providing the means for fast integration of information across areas with impact at both low and high frequencies. This may be especially important for gamma synchronization, where timing requirements are stringent (*18*). On the other hand, ephaptic interaction has been shown to enable the generation and propagation of slow waves in brain slices–even after they have been split (*5*). Similarly, SEFs could play a role in inter-hemispheric communication, bypassing corpus callosum connections. Other recent work suggests that they could play a role in the modulation of release of extracellular vesicles (*19*), a newly discovered form of cellular communication.

Relying on biophysical modeling and high-resolution neuroimaging analysis, we have built a first metric of ephaptic interaction in the human brain, characterizing its spatial distribution and its relationship with aging. Below we discuss the implications of ephaptic fields in humans, including their potential relevance for regulating brain oscillatory patterns and cortical excitability, their evolutionary meaning as well as potential role in neurological and pathological disorders.

### 3.1 Insights from models

Modeling results confirm many of the assumptions of the theoretical predictions. On the one hand, the decay of the electric field created by single dipole sources is confirmed to be well approximated by a 1/*r*^3^ power law, even in models that consider tissue heterogeneity. In the realistic model, multiple dipole sources create a field that decays slower (1/r^2^), as predicted by the 3D simplified sulcus model. This confirms that ephaptic interactions are limited to regions that are located close to one another. In the case of sulci, this limits interactions either to the cells close to the source(s) along the same wall, or cells on the opposite sulcus wall. We note that if the cortical region of interest is undergoing synchronous oscillations in a given band, the ephaptic effects will be in phase for dipoles along the same wall, and antiphase on the opposite wall. In our models with dipole density of 1.0 nAm/mm^2^, and assuming that the threshold for interaction was 0.1 V/m, ephaptic effects on the opposite sulcus wall could only be observed in the 3D toy model when the sulcus width was of 1 mm or less, and in the realistic 3D model in portions of the post-central sulcus where its width was the smallest (about 1.4 mm). For comparison, in Chiang et al. (*5*), a separation greater than 0.4 mm in a cut hippocampus slice was sufficient to impede ephaptic wave propagation (see Table 1), which, together with other findings, supports our selection of an analysis threshold of 0.1 V/m.

Further evidence that the scaling of the sources in these models is realistic comes from the observation that the maximum electrostatic potential recorded at scalp level in the realistic head model varied between 16 and 32 μV, respectively for a dipole density of 0.5 and 1.0 nAm/mm^2^. Since these dipoles comprise a cortical area of 5.3 cm^2^, these results seem consistent with the rule of thumb that ~6cm^2^ of activated cortical area are needed to produce detectable EEG at scalp level (*7*).

### 3.2 Topography of Ephaptic Fields in the Human Brain

As we have seen, EMOD1 is related to other metrics such as gyrification and cortical thickness. The latter is hardly surprising, since cross-sulcal ephaptic interaction requires the presence of cortical folding. The current study may provide further clues into the importance of gyrification as a zero-order proxy for ephaptic interaction. Studies have indicated that cortical gyrification is strongly and positively related to cortical volume but negatively related to cortical thickness in many regions of the cortex, and that frontal gyrification is positively related to performance in working memory and mental flexibility tasks (*20, 21*). Such results support the view that greater cortical gyrification is related to bigger brain volumes and better cognitive function. One advantage of gyrification is thought to be increased speed of brain cell communication, since cortical folds allow for cells to be closer to one other, requiring less time and energy to transmit neuronal electrical impulses (*17*). Ephaptic interactions and EMOD1 reflect similar advantages.

From an evolutionary point of view, we may hypothesize that natural selection forces that promoted folding the cortex to fit a larger cortical surface in a more static cranium (i.e., cortical gyrification), as a byproduct made available ephaptic interaction as a form of information transfer, which then also underwent natural selection. Across species, the degree of cortical folding correlates with brain weight and, more specifically, with cortical surface area. In all major mammalian lineages, the species with large brains tend to have more highly folded cortices than species with smaller brains (v. (*22*) and references therein). The pilot whale and bottlenose dolphin display the highest gyrification index values. The human brain, while larger than that of a horse, shows a similar gyrification index. Rodents generally display the lowest gyrification. Nonetheless, some rodents show gyrencephaly and a few primate species are quite lissencephalic. Research on the evolutionary biology studying ephaptic transmission is deeply needed.

### 3.3 Ephaptic Fields and Age

Analysis of the metrics computed on the MRI dataset indicate a robust correlation of EMOD1, cortical thickness, LGI, and surface area with age, as displayed in Figure 5. Not surprisingly, these metrics display moderate inter-correlations stemming from the covariation of cortical folding and sulcal separation. The index proposed here, which stems from physiological considerations related to ephaptic coupling, relies strongly on the notion of sulcal width and dipole strength (cubic) decay with distance. Studies of sulcal widening have shown it is associated to aging, decreased cognitive ability, dementia and schizophrenia (*15*). The negative association observed between EMOD1 and age suggest a highly speculative yet interesting scenario, where the decrease of ephaptic coupling with age may contribute to loss of control over oscillatory patterns and cortical excitability, potentially contributing to age-related cognitive changes. Furthermore, pathologies associated with cortical atrophy, e.g., dementia or traumatic brain injury, would alter ephaptic transmission as well, contributing to the pathophysiology as well as cognitive and behavioral symptoms.

Related to age-related changes in brain structures, the concept of “brain age” has been recently explored by multiple groups, looking at how structural MRI data can be used to estimate the “actual” biological age of a given brain as compared to his chronological age (*23–25*). Such analysis is carried out by fitting a model estimating chronological age by means of structural MRI data in a sample of age matched participants, to then compare residual values for each participant and label each brain as respectively “older” or “younger” than its reference cohort. Interestingly, estimated brain age has been shown to correlate with mortality, making a very interesting novel health biomarker (*24*). The structural properties such as LGI, thickness and grey matter density are considered, but no studies have investigated the potential role of ephaptic coupling distribution in determining brain age. Together with other potential mechanisms, such as functional reallocation of fMRI connectivity patterns, ephaptic coupling might constitute another key element to determine and maintain brain age.

### 3.4 Ephaptic role in neurological disorders

Hypersynchronized activity in seizure can generate large rhythmic fields of 20–70 V/m in the hippocampus and 3–9 V/m in the neocortex (v. (*26*)). Interictal discharges generate strong ephaptic perturbations that might very rapidly alter brain dynamics and cause, or at least contribute to, their deleterious effects on brain function and cognition, as also discussed in (*3*). Interestingly, cortical malformations of various types, including shallow sulci and defects of cellular migration, have been described in epilepsy as well (*27*), possibly linking cortical morphology and aberrant epileptic activity through alterations of ephaptic transmission.

More specifically, ephaptic interaction might play a role in the pathogenesis of seizure via its potential contribution to self-regulation of cortical excitability. As the cortical walls come in close proximity due to cortical folding, by projecting activity with the opposite phase on neighboring areas, ephaptic interaction might protect the brain from hypersynchronization. By the same token, the increasing amplitude and spatial extent of electrical activity generated during the last stage of a seizure (see, e.g. (*28*)) may act, through ephaptic interaction, as a homeostatic mechanism to end the seizure. Interestingly, focal cortical dysplasia lesions associated with epileptiform activity are preferentially located at the bottom of abnormally deep sulci (*29*), where such ephaptic homeostatic control would be weakest for geometric reasons.

Alteration of ephaptic interaction can also shed new light on other human brain disorders that are accompanied by change in cortical gyrification. For instance, Lissencephaly is a rare, genetically related brain malformation characterized by the absence of normal convolutions in the cerebral cortex and an abnormally small head. Symptoms may include unusual facial appearance, difficulty swallowing, failure to thrive, muscle spasms, seizures, and severe psychomotor retardation. Laminar heterotopia is a rare condition consisting in an extra layer of gray matter underlying properly migrated cortex, usually associated with epileptiform activity, cognitive deficits and alterations of functional connectivity patterns (*30,31*). Polymicrogyria is a condition in which the brain has an overly convoluted cortex. Symptoms can include seizures, delayed development or weakened muscles. Higher levels of gyrification are also found to relate to greater local connectivity in the brains of individuals with autism spectrum disorders, suggesting ephaptically mediated hyperconnectivity (*32*). The same could be predicted of healthy populations: increased ephaptic coupling (LGI and EMOD) would be associated to increased functionally connectivity, especially at high frequencies. Similarly, the brains of patients with schizophrenia also show reduced cortical thickness and increased gyrification when compared to healthy brains (*33*). Further studies on ephaptic transmission in various pathologies may offer novel insights to account for the identified alterations in brain oscillations and explain cognitive and behavioral symptomatology.

### 3.5 Relationship of tCS and ephaptic fields

Together with in-vitro and animal work demonstrating the physiological effects of weak electrical perturbations, abundant work in recent years indicates that weak electric fields applied over relatively large areas and over a duration of minutes can have significant physiological after-effects in humans (*34*). Interestingly, as highlighted above ephaptic fields are of the same order of magnitude as those generated by tCS, and both display large correlation scales (of the order of centimeters). In addition, in both types of electric fields are present in the cortex for relatively long times (minutes in tCS and indefinitely in ephaptic fields), and, at the scales of interest, at relatively low frequencies (≪ 1 kHz). These similarities suggest that the neuromodulatory effects of tCS may rely on a natural brain interaction mechanism.

For example, it is likely that the effects of tCS, which generates electric fields of the order of 0.1–2 V/m (as predicted by models and verified experimentally (*35, 36*)) may ultimately be explained by “spatiotemporal coherence” mechanisms, that is, to the augmented impact of weak but spatially extended, temporally coherent (DC or AC) and persistent (minutes) electric fields (*10, 37*) on neuronal networks in the presence of background noise. Such “array enhanced” emission and reception features would apply to both exogenous and endogenous fields.

A consequent question is how we can use these insights for better design of tCS protocols. If tCS leverages a natural and physiologically relevant ephaptic mechanism, understanding it in detail should provide valuable inputs for the design of optimized tCS in disorders such as epilepsy, depression or neuropathic pain, where questions remain on where to apply electric fields, for how long and with what temporal waveforms (DC, AC or endogenous, e.g., as derived from EEG), or, perhaps, to help understand what distinguishes treatment responders from non-responders. In particular, the design of tCS protocols should be conceived from the point of view of generating a summation of endogenous and exogeneous fields which the cortex will interact with as an endogenous one. For example, if age or atrophy (e.g., in dementia) predict a reduced impact of ephaptic interactions, would this also suggest a decrease of response to tCS? The hypothesis here would be that a brain that has lost the ability to engage in ephaptic communication will similarly be less sensitive to the effects of exogenous fields.

### 3.6 Limitations of the study and future directions

The conclusions drawn from our electric field models are subject to uncertainties in some parameters that may affect the volume conduction effects of the currents induced by the dipole sources. Some of these parameters are the conductivity properties of the tissues in the head in the low-frequency range of EEG. These conductivity values are known to considerably influence the electric field distribution in the brain, but the reported range of values in the literature is still somewhat inconsistent (*38*). They are also known to vary with individual anatomy, age and disease (*39–42*). Other important parameters in the model are dipole density and patch size. These are of critical importance, since they influence the location and size of the areas which are influenced by source activity.

Another important limitation in this study is the use of a simplified metric (EMOD1) as opposed to a full calculation of the ephaptic field generated by cortical dipoles (EMOD proper, Eq. 4). This represents a convenient trade-off to be able to evaluate this metric on a large dataset, but it may be improved in the future. In addition, we have used here an interaction model which does not consider the complexity or spatial distribution of pyramidal neurons, or the effects on other types of neurons, much as it is done in practice, with some justification (*10, 43*)(*10, 33*), in the analysis of tCS effects. Finally, the effects of tCS have been studied in computational models of the brain (*43–45*) using the lambda-E model discussed above and ignoring the intricacies of cortical network circuitry. This is a simple model that will be improved in the future.

Further work remains to be carried out to disentangle the differential contributions of EMOD1, cortical thickness and other cortical morphologies to explaining measures of brain function and cognition. An interesting line of research will be to determine computationally the impact of ephaptic fields on neuronal dynamics in both the healthy and pathological cortex, along the lines of (*45*).

## 4 Conclusions

Our findings, in line with earlier experimental work, suggest that ephaptic transmission could constitute the basis of a novel framework for the understanding of brain function and human cognition, as well as neurological and psychiatric pathology where brain structural alterations are present.

## 5 MATERIALS AND METHODS

### 5.1 Mechanisms

Given their anatomical characteristics (elongated form factor, which enhances the effects of electric field on membrane polarization), organization (horizontal connectivity, homogeneous orientation in cortical patches and temporal synchrony (*8*)), cortical pyramidal cells are well suited as electric field generators (*8*). In analogy with reciprocity principles that apply to electromagnetic radiation antennae, for the same reason they are good field sensors of quasi-static (endogenous or exogenous) electric fields. Other cortical neuron types, however, may also play a role.

tCS (also known as tES) is a family of noninvasive techniques that include direct current (tDCS), alternating current (tACS), random noise current stimulation (tRNS) or others using specially designed waveforms. It consists in the delivery of weak current waveforms through the scalp (with electrode current intensity to area ratios of about 0.3–5 A/m^2^) at low frequencies (0–1 kHz) resulting in weak but spatially extended electric fields in the brain (with amplitudes of about 0.1–2 V/m). tCS is applied during several minutes (typically ~20 minutes). Such electric fields do not initiate *per se* action potentials, but they can influence the likelihood of neuronal firing by the modulation of neuronal transmembrane potentials in relatively large cortical patches, resulting in changes in firing rates and spike timing. The sustained application of such weak fields during sufficiently long periods of time (several minutes) leads to plastic changes of neuronal connectivity through Hebbian mechanisms (see, e.g., (*46–48*)). Thus, like SEFs, the main characteristics of exogenous tCS macroscopic fields are that they are weak, low frequency with moderate to large spatial correlation scales (> 1 cm), and, in practice, applied for relatively long times.

The concurrent effects of tCS are understood to be mediated by the coupling of electric fields to ordered populations of elongated neurons, especially pyramidal cells (see (*10, 49*) and references therein). Neurons are influenced mostly by the component of the electric field parallel to their trajectory (*2, 50–53*), and, therefore, knowledge about the orientation of the electric field is crucial to predict the effects of stimulation. The components of the field perpendicular and parallel to the cortical surface are of special importance since pyramidal cells near the cortical surface are mostly aligned perpendicularly to the surface, while many cortical interneurons and axonal projections of pyramidal cells tend to align parallel to the surface (*54–56*). For a long, straight finite fiber with space constant *λ* in a homogeneous electric vector field E, the transmembrane potential difference is largest at the fiber termination, with a value that can be approximated to first order by

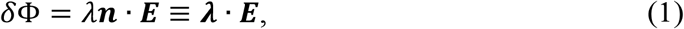

where ***n*** is the unit vector defining the fiber axis in the orthodromic direction (see Figure 1). In this approximation, which is sometimes called the “lambda-E model” (*10, 57*), the spatial scale is provided by the neuron space constant, and the effect is modulated by the relative orientation of field and elongated neuronal populations. The effect is thus determined by both field magnitude and by its direction.

Although membrane perturbations from weak fields are sub-threshold (about 0.1–0.2 mV per V/m applied (*49*)—significantly lower than the 20 mV depolarization required to bring a neuron from resting potential to spike threshold in vitro (*58*))—, nonlinear effects in coupled populations probably lead to an amplification of effects. For example, mathematical models have demonstrated the amplification of weak but coherent signals in networks of nonlinear oscillators (see, e.g., (*49–51*)(*59, 59, 60*)) and, more specifically, in computational models of neural circuits (*2, 3*)). This effect is ultimately dependent on the coupling strength of network elements and their architecture, while noise can contribute to the enhancement of small but homogeneous perturbations in the network (array enhanced stochastic resonance). Thus, co-operative effects arising from noise and coupling in coupled systems can lead to an enhancement of the network response over that of a single element. Such amplification mechanisms could also play a role in other phenomena where a surprising sensitivity to weak perturbations has been found, as with the effects of Earth-strength magnetic field rotations in EEG alpha band activity (*61*).

In summary, assemblies of neurons, if appropriately and homogenously oriented, can function as antennae for ephaptic coupling. We adopt here the lambda-E model to estimate ephaptic effects, given the similar features of exogenous and endogenous fields of interest.

### 5.2 Estimates of endogenous field strength from reciprocity arguments and EEG

While in the next sections we model SEFs in the cortex using finite element models, here we provide some estimates from reciprocity considerations (*62–64*) by leveraging earlier work modeling the electric fields generated by tCS. Realistic head modeling shows that tCS fields associated to typical 1 mA bipolar transcranial current injection montages are of the order of 0-0.5 V/m (electric field normal to the cortex, *E_n_*) (*65*), and about 5–10 times smaller when averaged over cortical patches at tCS resolution scales (several cm^2^). These models have now been validated by invasive measurements (*6, 35, 36*), where a bipolar current of about 1 mA leads to median electric field magnitudes of the order of 0.1 V/m.

According to the reciprocity theorem, the magnitude of the E-field normal to the cortical surface induced by a given tCS montage is proportional to the sensitivity of the same montage when used for EEG to monitor the electrical signals generated by a dipole source at the same point in the cortical surface and oriented perpendicularly to it. Let us denote by *V_ab_* = 1 *mA* the current applied from point *a* to point *b* in the scalp that induces the cortically normal electric field *E_n_* somewhere at a point *x* in the cortex. Consider a hypothetical reciprocal EEG measurement where we observe a potential difference *V_ab_* = 10 *μV* between the same points *a* and *b* produced by a dipole located at *x* and normal to the cortical surface—such as the one in Figure 1. The reciprocity theorem implies that we can replace the pair (*E_n_, I_ab_*) with (*V_ab_, p*) with the ratio of the first pair the same as the ratio of the second. Hence, from the current-electric field data pair we can deduce, given *V_ab_*, a value for a reciprocal dipole *p: V_ab_/p* = −*E_n_/I_ab_*, which implies |*p*| = |*I_ab_V_ab_/E_n_*| = 10 x 10^−6^ *V* x 10^−3^ *A*/(0.1 *V/m*) = 100 *nA · m* using a value of *E_n_* ≈ 0.1 *V/m*. So, ifa lone dipole located at *x* were responsible for the observed *V_ab_*, it would have this strength. As an example, for the chosen realistic head model sulcus model described below (Section 5.4), we calculated the voltage distribution at the scalp induced by such a single 100 nAm source dipole. The dipole was oriented normal to the cortex. At that location, the normal component of the electric field generated by a montage with CP2 as the anode (1 mA) and T10 as the cathode (−1 mA) was of 0.13 V/m. The potential difference between electrodes CP2 and T10 was of 13.1 μV, in agreement with the reciprocity calculation.

Given such a dipole ***p*** at location *x*, what is the associated ***E*** at some nearby point *y*? As a first approximation, the electric field from a current dipole in a homogeneous conductive medium is (in polar coordinates, see (*64*), p. 33):

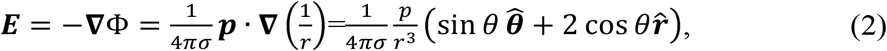

where *r* is the distance between *x* and *y*, and *σ* the conductivity of the medium. For example, the field magnitude at 1 mm of distance from the idealized dipole on the contiguous cortical surface is *E* ≈ 40 *V/m* (*θ* = 0, *σ* = 0.40 *S/m* in grey matter tissue, see for instance (*66*)). This is at the high end of DC stimulation regime experiments (in-vitro, see Table 2). At 1 cm distance from the dipole, *E* = 0.05 *V/m*. Out in the CSF, where (*θ* = 90, *σ* = 1.79 *S/m*), the magnitudes are *E* ≈ 4 and 0.004 *V/m*, respectively. The dipole approximation is applicable for distances significantly larger than the dipole size (the space constant of pyramidal neurons is typically much less than 1 mm, see e.g. (*67*)).

Of course, EEG signals are not generated by single point dipoles but by the summation of fields from extended sources (coherent patches) and collections of them. Despite of this, to the extent that these sources are small compared to scales we are interested in, these estimates give an order of magnitude of what we may expect to observe. Measurements in the human neocortex indicate that current dipole surface densities in the cortex are in the range of 0.16–0.77 *nA · m/mm*^2^ (*16, 17*). There appears to be a maximum value across brain structures and species (1–2 *nA · m/mm*^2^). Studies using combined electrocorticography and MEG show that coherent area sizes of the order of 1 to 20 *cm*^2^ are needed for MEG detection, with the larger ones observed in epileptic discharges (*68*). At a density of 0.25 *nA · m/mm*^2^, our hypothetical dipole of 100 *nA · m* above would be realized over a patch of about 4 *cm*^2^.

Finally, we note that cortical folds bring together pyramidal populations of opposite orientation to distances of much less than 1 *cm* (even submillimeter) which should play an important role in extending the effects of dipole fields beyond their immediate neighborhoods.

### 5.3 Simplified 3D volume conductor model of ephaptic interactions

To investigate in more detail the electric field distribution created by dipole sources on a heterogeneous volume conductor, we first created a 3D finite element toy model. The model, shown in Figure 2, includes a simplified representation of a sulcus and of the scalp, skull, cerebrospinal fluid (CSF), grey-matter (GM) and white-matter (WM) tissues. This geometry was then extruded 100 *mm* along the z-axis (out of plane direction). Sources were placed in a patch located in the posterior wall of the sulcus, in the GM-CSF interface.

**Figure 2:**
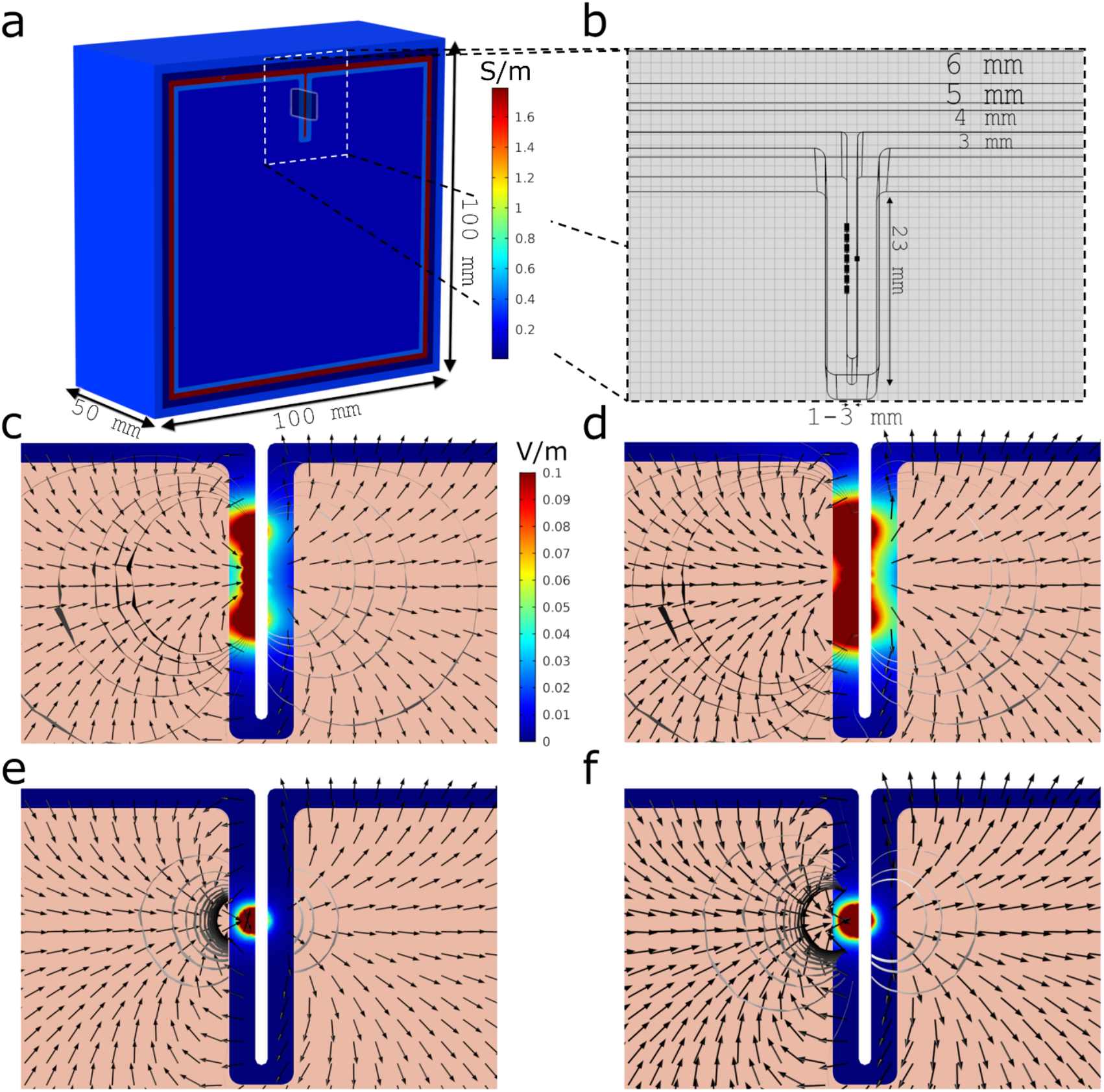
Geometry and electric field distribution in 2D model of a sulcus. (a) 3D view of half of the simplified volume conductor (100×100×100 mm). The different tissues are colored by their respective conductivity, in S/m. The patch of single dipole sources is placed in the central region of the model (posterior wall of the sulcus), covering an area of 60 mm^2^. (b) Sagittal view of the model (sulcus width of 1 mm) with dipole sources in its posterior wall. (c-f) Magnitude of the electric field in the GM tissue for models with different source strength and patch distributions (common color scale between plots in V/m). Also shown are vector plots of the electric field and isosurfaces of the electrostatic potential. Left/right columns represent the models with the sources scaled to a density of 0.5 and 1.0 nAm/mm^2^ respectively. Top/bottom rows represent multiple/single dipole distributions.

The tissues were assumed to be homogeneous and isotropic, with electrical conductivity values appropriate to the low frequency range of interest (*65, 66*): 0.33,0.008,1.79,0.40 and 0.15 *S/m* respectively for scalp, skull, CSF, GM and WM. Sources were modeled as point dipoles, with a direction perpendicular to the sulcus wall. Two models for the sources were built: a single dipole model and a multiple dipole model (77 dipoles located in a 1 *mm* × 1 *mm* regular grid comprising a 60 *mm*^2^ patch — as shown in Figure 2). The single dipole model was used to study the electric field distribution of a dipole source and its decay with distance. The multiple dipole model was used as a more realistic representation of a patch of sources. For each source model, the sulcus width was varied between 1 and 3 mm, which are median sulcus width values on the low/high-end of the reported sulci width for subjects between 20 and 80 years of age (*15*). All models were solved in Comsol with the AC/DC package (v5.3a, www.comsol.com). The finite element mesh comprised tetrahedral second order Lagrange elements with a minimum size in the GM and CSF layers of 0.5 mm. Dipole sources were modeled with Comsol’s “Electric Point Dipole” boundary condition, which allows the user to specify the direction and strength of the dipole.

### 5.4 Building a realistic brain model of ephaptic fields

The electric fields generated in the brain with tCS can now be readily modeled at the individual level using imaging data (see (*57, 69*) for recent reviews). We employ here the same techniques to model endogenous fields from cortical dipoles, that is, finite element modeling derived from MRI (see Figure 3). The model, described in detail in (*65*), is based on the Colin27 MRI dataset (http://www.bic.mni.mcgill.ca/ServicesAtlases/Colin27). It includes realistic representations of the scalp, skull, CSF (including ventricles), GM and WM. Each tissue was modeled as explained in the previous section. Dipole sources were placed in the grey matter-cerebrospinal fluid (GM-CSF) surface of the model, perpendicularly to it, in similar fashion to what was done in the 3D simplified model. As before, two source distributions were calculated: a single node source mode and a multiple source model comprising a cortical surface of 5.30 cm^2^. In the single source model, the cortical surface was parcellated into 112 AAL areas and a point was chosen randomly in each area, for a total of 112 single source models. The multiple source model was built by placing 133 dipole nodes in the posterior wall of the post-central sulcus (see Figure 3 b). All electric field calculations were performed in *Comsol* with the AC/DC package.

**Figure 3:**
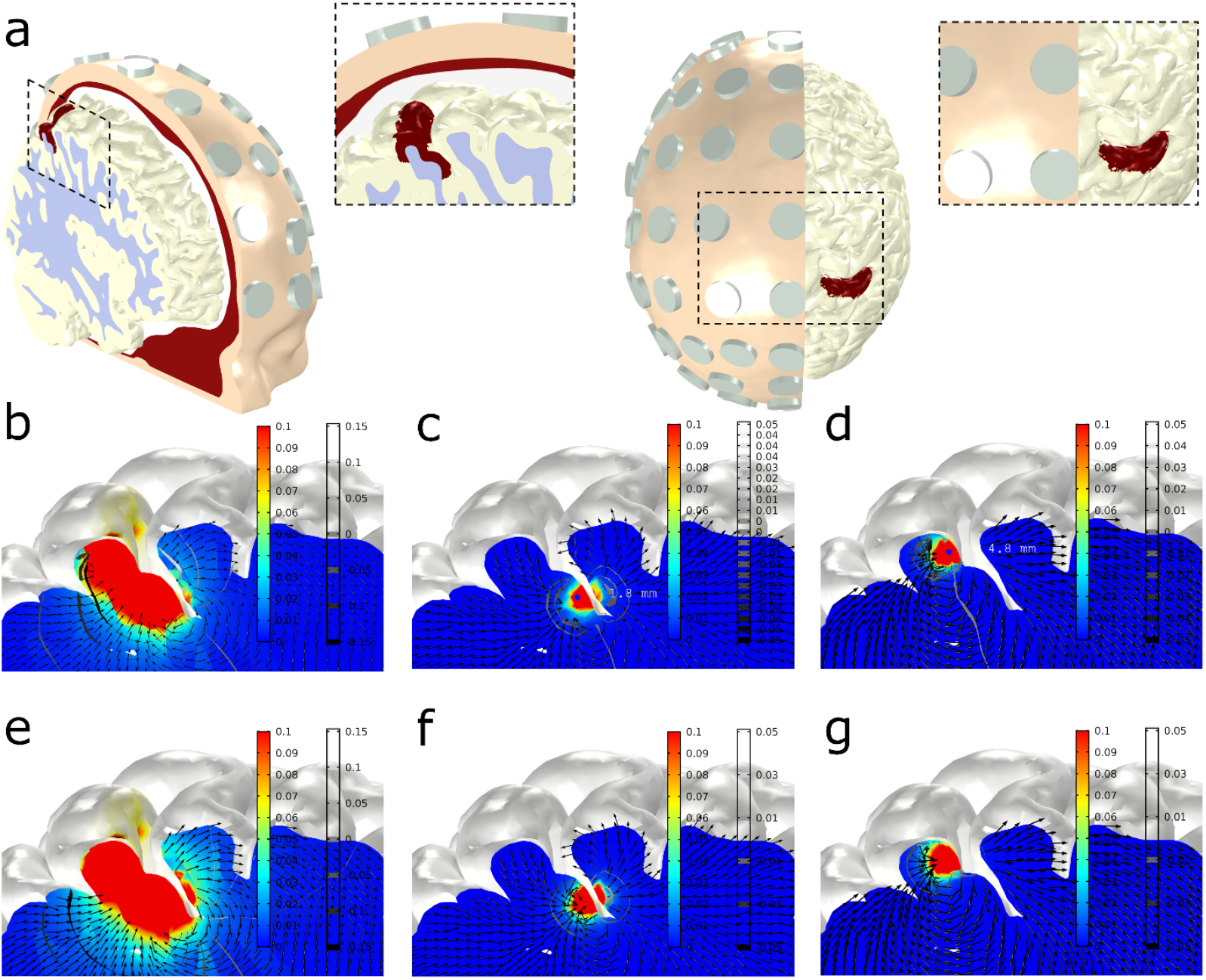
Realistic head model. **(a)** Two views of the 3D volume conductor geometry, including volumes representing the scalp (yellow), skull (red), CSF (white), GM (grey) and WM (light red). Models of electrodes, placed in the 10-10 EEG positions, are also included in the model (grey). The patch used to place dipoles in the multiple-source model (posterior wall of the post-central sulcus, on the right hemisphere) is displayed in red in the GM volume. It comprises a cortical surface of 5.30 cm^2^. The captions provide zoomed views of the cortical patch with the dipole sources. **(b-g):** Electric field magnitude (color bar in V/m) and vector field direction, and isosurfaces of the electrostatic potential (gray-scale, mV) in a sagittal slice passing through the middle of the right hemisphere post-central sulcus. First (b-d) and second (e-f) rows: dipole density per unit area of 0.5/1.0 nAm/mm^2^. Columns, from left to right: model with all dipole sources, model with single dipole in narrow region of the sulcus, model with single dipole in wide region of the sulcus. The location of the individual dipoles in the middle and right-most columns are shown as blue circles in figures c and d. The sulcus is approximately 5.5 mm wide in its wide region and 1.8 mm wide in its narrow region.

**Figure 4:**
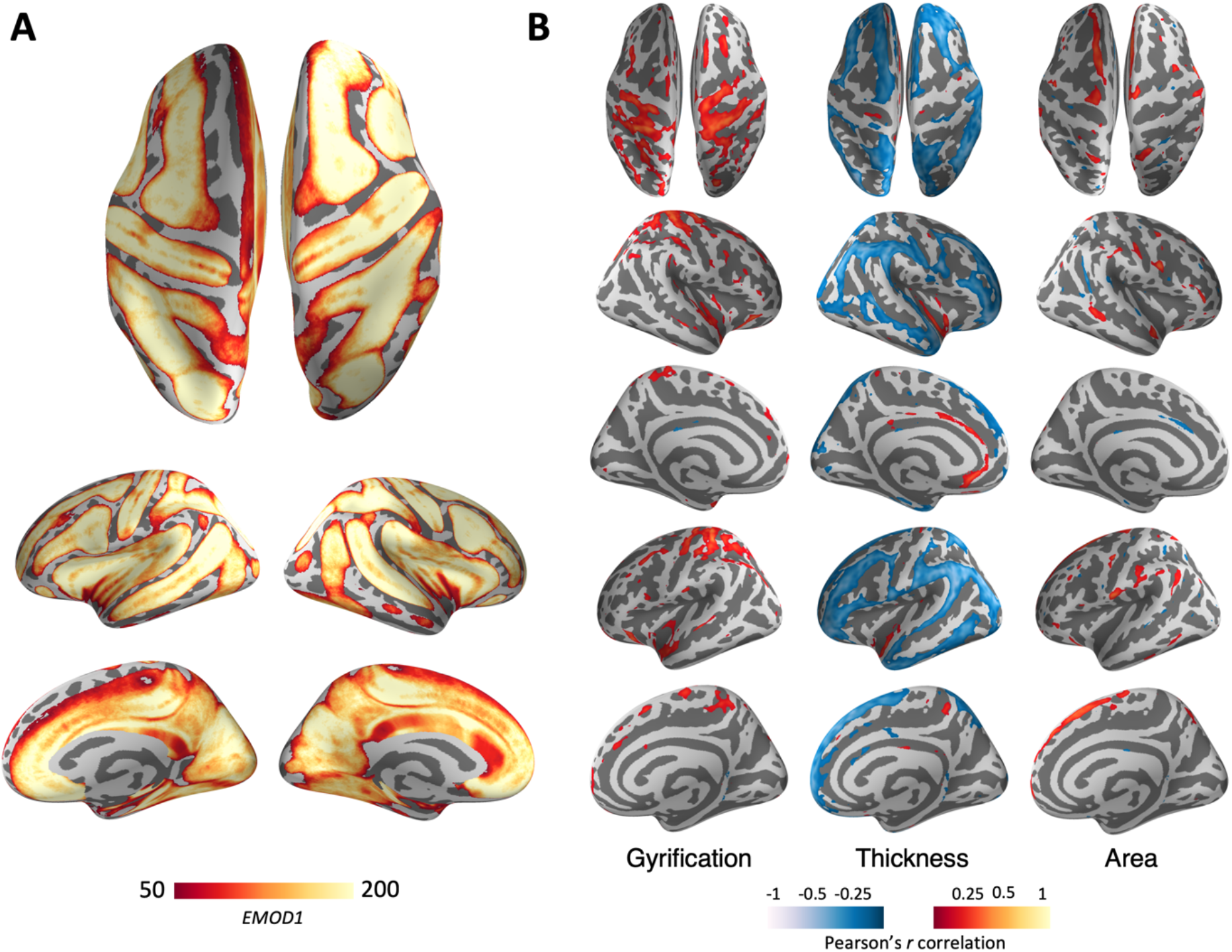
Ephaptic Modulation in the human brain. **(A) Average EMOD1.** Individual EMOD1 maps are registered to Freesurfer’s common template (*fsaverage*) and then averaged at each vertex across subjects. For the purpose of visualization, we have thresholded the average EMOD1 map at EMOD1>50. **(B) Vertex-wise correlation.** At each vertex, the Pearson’s correlation coefficient between EMOD1 and cortical surface area, thickness, gyrification and subject’s age is computed. The resulting maps are then corrected for multiple comparisons using the Benjamini-Hochberg procedure (p-value <0.05). Pearson’s correlation coefficient values for vertices that passed the multiple comparison correction are overlaid on Freesrufer common template *fsaverage*).

**Figure 5.**
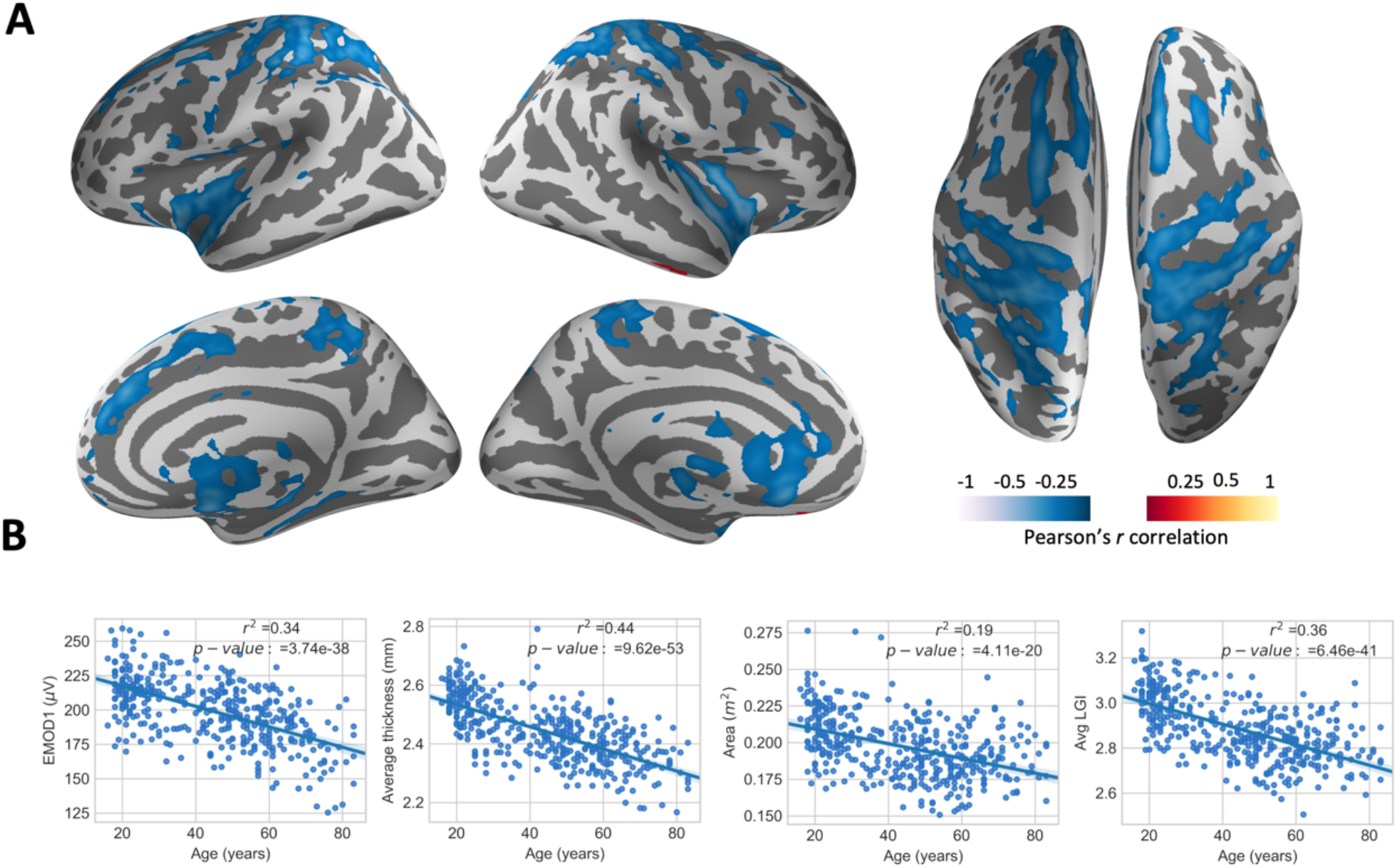
Ephaptic interaction and Aging EMOD1. (A) Vertex-wise EMOD1 values were correlated with age across the sample of 401 subjects, resulting in a weighted map displaying the cortical regions whose ephaptic modulation index is significantly affected by aging. (B) Individual data for correlation between age, EMOD1, as well as cortical morphologies are displayed. Red-yellow shows positive and blue-cyan negative correlations.

### 5.5 Ephaptic modulation index (EMOD and EMOD1)

In this section we define an index to estimate, for a given individual brain model, the role of ephaptic modulation. The index provides an average over the cortex of the impact that emitting dipoles have on receivers. We have considered several aspects to define it meaningfully. First, it should reflect the basic physics of dipoles (field decay with distance) and coupling to neurons (directional lambda-E model (*10*)). Second, it should be insensitive to local effects of a dipole on its local neighbors on the cortical manifold, as this will be a strong but unspecific effect. Rather, it should emphasize the effects of neighboring dipoles across-sulcus. Finally, for ephaptic effects from near dipoles to add to some relevant value, they should be *coherent* in time. This means the metric should disregard remote sources (e.g., a few cm away), which will be presumably less coherent. The coherence space scale in the cortex depends on the frequency of the dynamics of interest. For instance, the spatial correlation length of dipole activity in the cortex is larger at lower frequencies. It is often stated that a coherent patch of 6 cm^2^ is needed to create signals that can be detected by EEG (*7*). It is for these reasons that EEG power is weaker at high frequencies (there is no frequency dependence on conductivity at the frequencies of interest, as discussed in (*70*)). This also indicates that ephaptic effects are probably frequency dependent, and stronger at low frequencies.

Now, using the lambda-E tCS interaction model, the ephaptic impact of a source dipole at y on a neuron or neuron population receiver at x (in μ*V*) may be approximated by *ε_y_*(*x*) = ***λ_x_ · E_y_***(*x*), where ***E_y_***(*x*) is the endogenous electric field vector at *x* generated by a dipole at *y* and ***λ_x_*** the space constant vector of the receiver neuron or neuronal population at x. The membrane perturbation may be positive (depolarizing) or negative (hyperpolarizing).

We sum ephaptic the contributions from dipole generators over the cortical mesh surface (all *y* ≠ *x*) *to* produce a total ephaptic impact factor for each cortical location *x* is (in *μ*V),

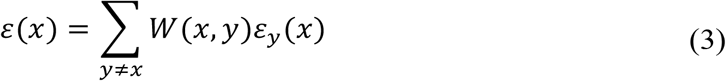

where *W* (*x, y*) is a support function to account for the requirements of non-local but coherent (not too far) contributions. This is a local measure on the cortical surface, which we can use to produce cortical surface maps of ephaptic effects.

In the same vein, the average global index equation for a cortex is simply (μ*V*):

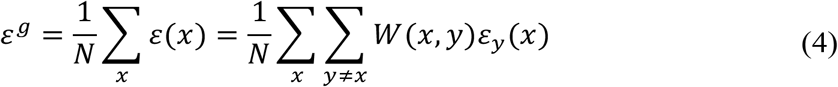

with N the number of nodes in the cortical mesh.

While Equation 4 provides a generic, precise expression (EMOD), it may be hard to compute in practice (a realistic head model of cortical dipole electric field at each node needs to be evaluated). We may approximate it using Equation 2 for very short distances and mutually opposed emitter/receiver dipoles (with *θ* = 0) as

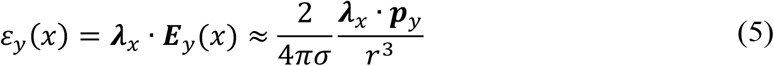

We will set ***p***_*y*_ = *p*_0_*δ****An***_*y*_ with *p*_0_ = 0.5*nA · m/mm*^2^ and ***λ***_*x*_ = *λ*_0_***n***_*x*_ with *λ*_0_ = 1 *mm*. We denote the local unit normal vector at the source at *y* by ***n***_*y*_. We collect some of these factors into a constant for use below, *κ* = *λ*_0_*p*_0_/(2*πσ*) (with conductivity evaluated at GM). Based on this, we provide a simplified approximation which uses the fact that dipole strength falls, approximately, as the cube of the distance, with ***n***_*x*_ and ***n***_*y*_ denoting local unit cortical surface normal vectors at source and receiver locations,

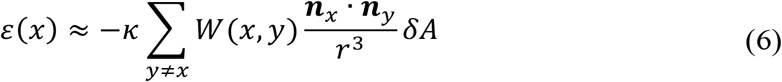

This index takes into account orientation of dipole and affected populations, and in particular, if the effect of the dipole on other regions is excitatory or inhibition. Finally, to select contributions from near dipoles in Euclidean space but geodesically distant on the surface (e.g., across sulci with opposed orientation), we write

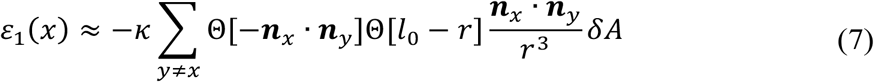

and

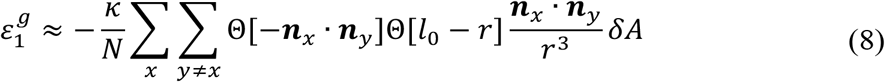

that is, with the weighting term *W* (*x, y*) = Θ[−***n***_*x*_ · ***n***_*y*_]Θ[*l*_0_ − *r*], with Θ[*x*] the Heaviside step function (defined as Θ[*x*] for *x* ≤ 0 and 1 otherwise) and *l*_0_ a scale relevant for interaction (maximal distance to consider coherent contributions). We set *l*_0_ = 5 *mm*.

We call this simplified index EMOD1 (see Supplementary Materials for a discussion on variants of EMOD1). It can be computed vertex-wise to produce cortical maps or averaged over the surface. Its calculation requires only the segmentation of the cortical surface and calculation of surface normal vectors from MRI images.

### 5.6 Imaging data and analysis

To test the variation of the ephaptic modulation index with age, we calculated it (using the simplified expression in Equations 7 and 8) for 401 subjects with ages between 16–83 years using a publicly available database. High-quality structural T1-weighted MRIs (3T) were acquired for 401 subjects from the NKI-Rockland database (*71*). MRI images were acquired using a 3-T Siemens MAGNETOM TrioTim with the following parameters: MPRAGE sequence, TR = 1900ms, TE =2.52ms, and TI=900ms, Flip Angle=9 degrees, FOV=250×250mm, voxel size=1 mm isotropic.

Structural T1-weighted MRIs were processed using the Freesurfer v6.0 software package to create three-dimensional representations of cortical surface (*72*). The Freesurfer pipeline includes automated Talairach transformation, segmentation of subcortical white matter and deep grey matter structures based on intensity and neighbor constraints, intensity normalization, tessellation of grey matter-white matter boundary and grey matter-CSF boundary, automated topology correction and reconstruction of cortical surface meshes (*73*). Next, reconstructed white surfaces were registered to Freesurfer template (*fsaverage*) based on cortical folding patterns using spherical registration implemented in Freesurfer (*mri surf2surf*).

For each subject, we also have computed cortical morphometrics including cortical thickness, surface area, and gyrification. Gyrification quantifies the cortical surface hidden in the sulci as compared to the visible cortical surface. The vertex-wise cortical gyrification was measured by calculating the gyrification index in circular three-dimensional regions of interest (*74*). This method uses an outer smooth surface tightly wrapping the pial surface and computes the ratio between areas of circular regions on the outer surface and their corresponding circular patches on the pial surface (see https://surfer.nmr.mgh.harvard.edu/fswiki/LGI for a description of how to calculate it with Freesurfer). At each vertex, cortical thickness was measured as the distance between white and pial surfaces, and cortical surface area was calculated by averaging the area of all faces that meet at a given vertex on the white matter surface.

Spherical registration implemented in Freesurfer (mri surf2surf) was used to register white matter surfaces into *Freesurfer* common template (*fsaverage*) to perform group-level analyses. We used 10 mm full-width-at-half-maximum (FWHM) Gaussian kernel to smooth cortical thickness, surface area, gyrification and EMOD1 maps.

### 5.7 EMOD calculation

For EMOD1 calculation, the GM meshes obtained from Freesurfer were corrected from morphological defects using the *Mayavi* (https://docs.enthought.com/mayavi/mayavi/) and *Pymeshfix* (https://pypi.org/project/pymeshfix/) toolboxes for *Python*. Surface normal vectors were then calculated in Matlab (v2018a, www.matlab.com) using the Iso2Mesh pipeline (http://iso2mesh.sourceforge.net/cgi-bin/index.cgi). For each mesh point of the surface we also calculated the Euclidean distances to all the other points in the mesh, and used this information to compute EMOD1 locally and then globally using Equations 7 and 8.

### 5.8 Statistical Analysis

Statistical analysis of correlations of metrics with age has been carried out using the Pearson correlation coefficient and its associated statistical significance using the Student’s t-distribution. All regressions were performed with the *Statsmodels* package for Python (*75*).

We performed vertex-wise Pearson’s correlation analyses between EMOD1 and cortical morphologies (cortical thickness, surface area and gyrification) as well as subjects’ age. False discovery rate (FDR) approach was used to control for multiple comparisons (Benjamini-Hochberg procedure, corrected p-value < 0.05) (*76*).

## Acknowledgements/Conflict of Interest

Giulio Ruffini is co-founder of Neuroelectrics; Ricardo Salvador and Roser Sanchez-Todo work at Neuroelectrics, a company that produces EEG and tCS systems. Giulio Ruffini would like to thank Niels Birbaumer for inspiring discussions on the relation of tCS with EEG neurofeedback. GR is partially supported by the European FET Open project Luminous (European Union’s Horizon 2020 research and innovation programme under grant agreement No 686764). Dr. Pascual-Leone and Dr. Santarnecchi are partially supported by Office of the Director of National Intelligence (ODNI), Intelligence Advanced Research Projects Activity (IARPA), via 2014-13121700007. The views and conclusions contained herein are those of the authors and should not be interpreted as necessarily representing the official policies or endorsements, either expressed or implied, of the ODNI, IARPA, or the U.S. Government. Pascual-Leone is further supported by the Berenson-Allen Foundation, the Sidney R. Baer Jr. Foundation, grants from the National Institutes of Health (R01HD069776, R01NS073601, R21 MH099196, R21 NS082870, R21 NS085491, R21 HD07616), and Harvard Catalyst | The Harvard Clinical and Translational Science Center (NCRR and the NCATS NIH, UL1 RR025758). Dr. Santarnecchi is supported by the Beth Israel Deaconess Medical Center (BIDMC) via the Chief Academic Officer (CAO) Award 2017, the Defence Advanced Research Projects Agency (DARPA) via HR001117S0030, and the NIH (P01 AG031720-06A1, R01 MH117063-01, R01 AG060981-01). The content of this paper is solely the responsibility of the authors and does not necessarily represent the official views of Harvard University and its affiliated academic health care centres, the National Institutes of Health, the Sidney R. Baer Jr. Foundation.

## SUPPLEMENTARY MATERIALS

### Speed of electromagnetic waves in the brain

Table S1 provides a summary of the speed of electromagnetic waves in brain media.

### Review of literature on the effects of slow, weak electric fields (SEFs)

See Table S2 for an overview of relevant papers involving weak fields.

### Decay of dipole fields

Figure S1 displays plots with the decay of electric field and potential as a function of Euclidean distance for different models.

### 3D sulcus geometry

Figure S2 displays distance measurements of the sulcus gap.

### EMOD1 maps for selected subjects

Figure S3 displays the surface distribution of the EMOD1 coefficient (l0 of 5 mm) for subjects with different ages.

### Variants of EMOD1

We provide here some variants of EMOD1. We recall the definition of EMOD1 (with *l*0 = 5 mm):

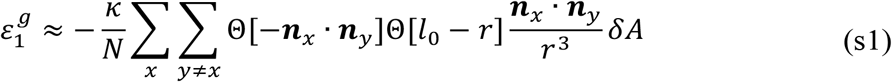

The spatial scale *l*_0_ can be varied, but it does not have a big impact on the results.

The first main EMOD1 variant just considers the effect of distance between emitter and receiver, ignoring relative orientation:

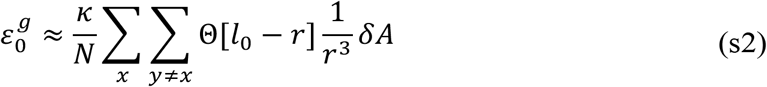

The second one takes into account relative orientation, but does not enforce the requirement in EMOD1 for opposite orientation of emitter and receiver (which forces cross-sulcal contributions in EMOD1):

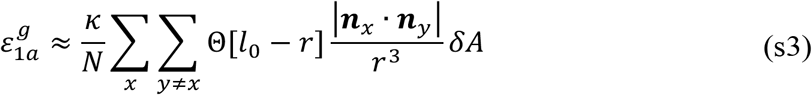

Figure S4 provides linear fits of EMOD variants with age, S5 provides second order fits.

### Second order correlations of metrics

Figure S6 provides second order fits of EMOD1, LGI, cortical thickness and area with to age, while figures S7 and S8 provide Pearson cross-correlation between the different metrics.

### Scalp map/EEG generated by dipole patch model

Figure S9 displays the scalp map potential for one of the chosen dipole cortical patches (Figure 3, 0.5 nAm/mm^2^ density).

**Table S1:** Relative permittivity, speed of light reduction factor with respect to vacuum (*c/v*), speed of light in tissue (v) in the low frequency range (10 Hz) for various tissues, with data from (*77*) provided online at http://niremf.ifac.cnr.it/tissprop/. Here we use 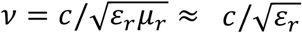 (the relative magnetic permittivity in body tissues is close to unity (*78*)). The last column is the time in nanoseconds required by ephaptic signals to traverse a sphere of 20 cm. Speed increases 3–4 times at 100 Hz for grey matter (GM) and white matter (WM), and stays constant for cerebrospinal fluid (CSF).

| Tissue | $\epsilon_r$ | $c/v$ | $v$ (km/s) | $\tau_{20\text{ cm}}$ (ns) |
| --- | --- | --- | --- | --- |
| Vacuum | 1 | 1 | 299,792 | 0.0 |
| CSF | 109 | 10 | 28,715 | 0.0 |
| GM | 40,699,000 | 6,380 | 47 | 4.3 |
| WM | 27,627,000 | 5,256 | 57 | 3.5 |

**Table S2:**
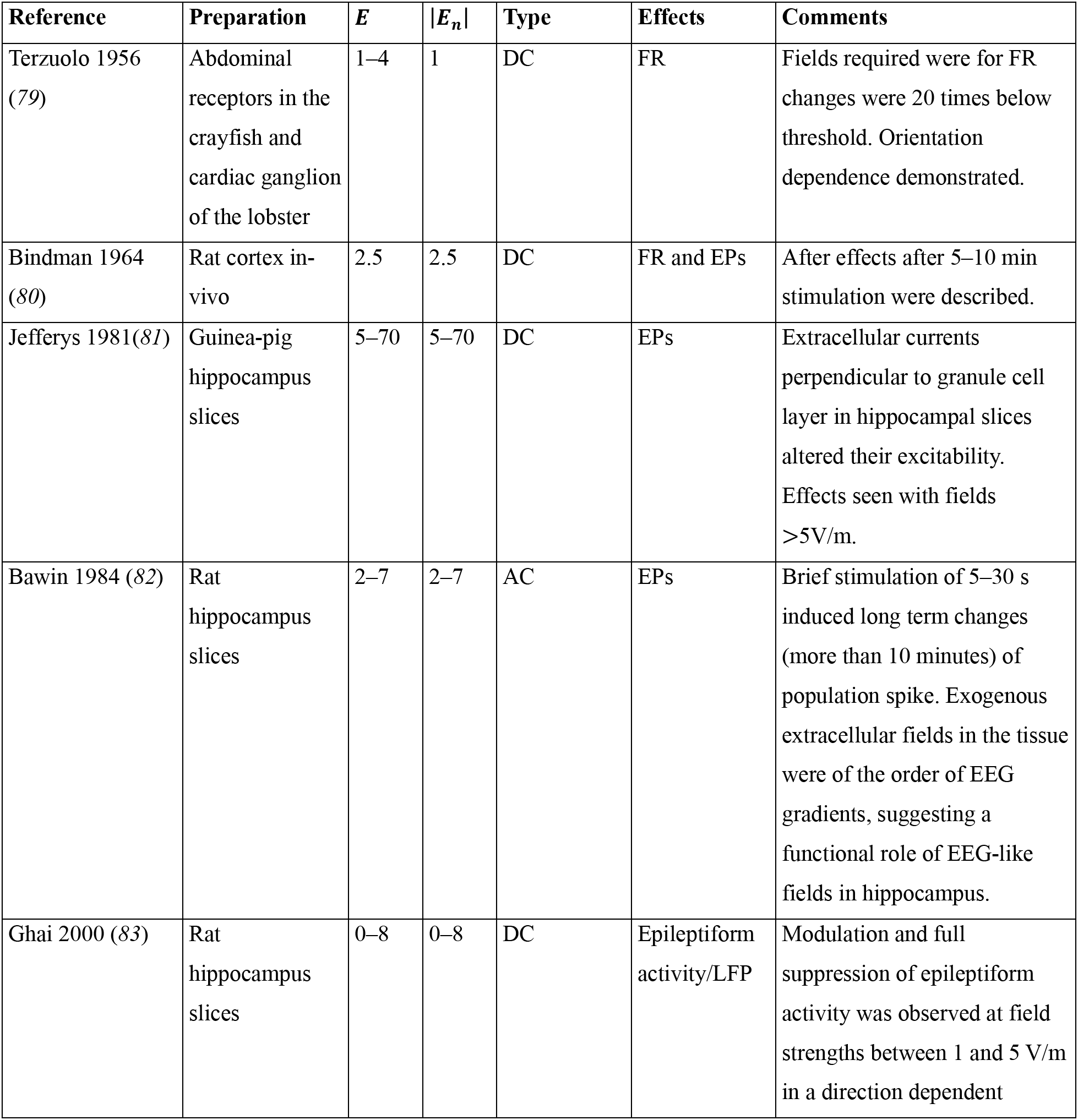
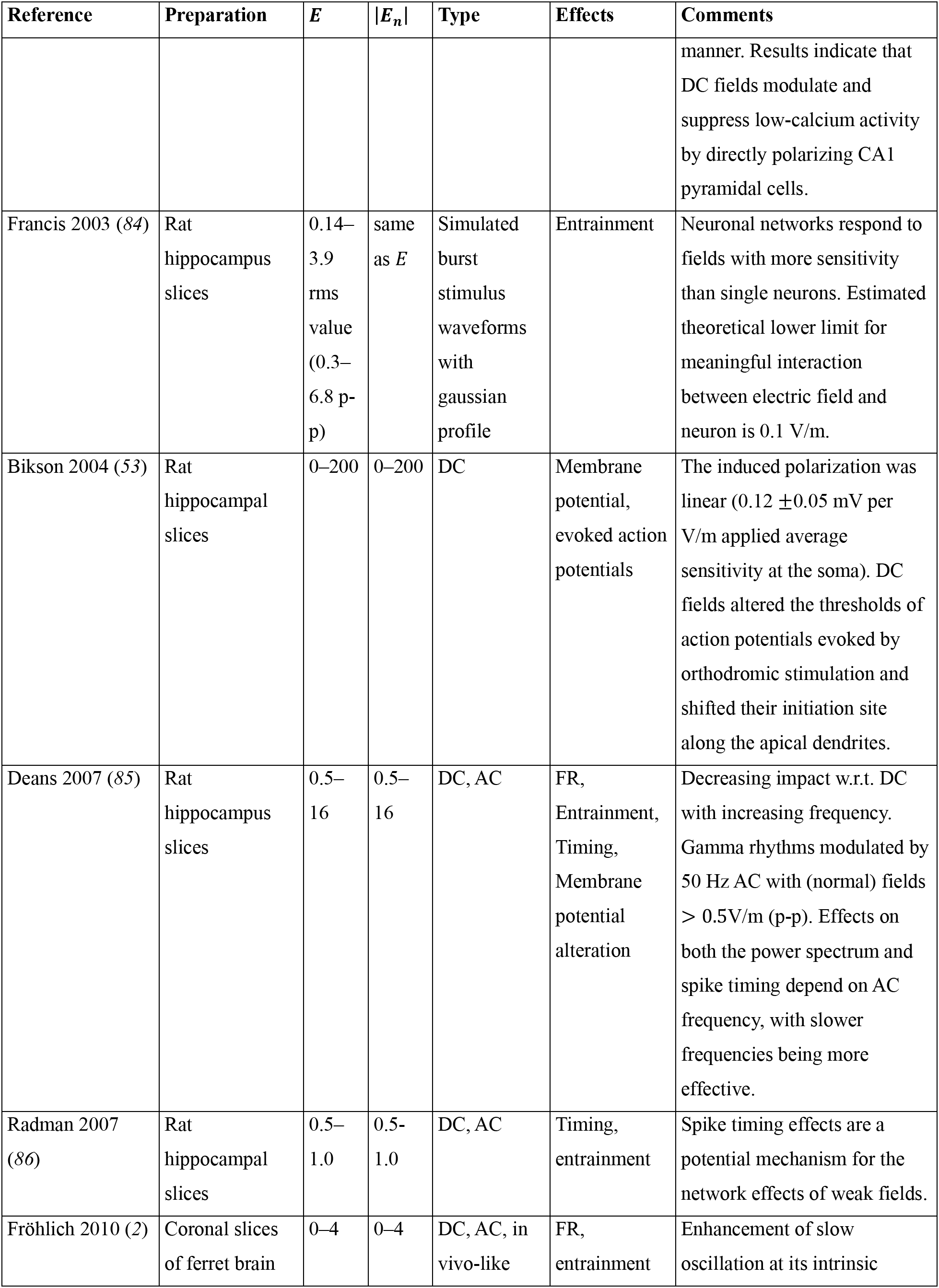
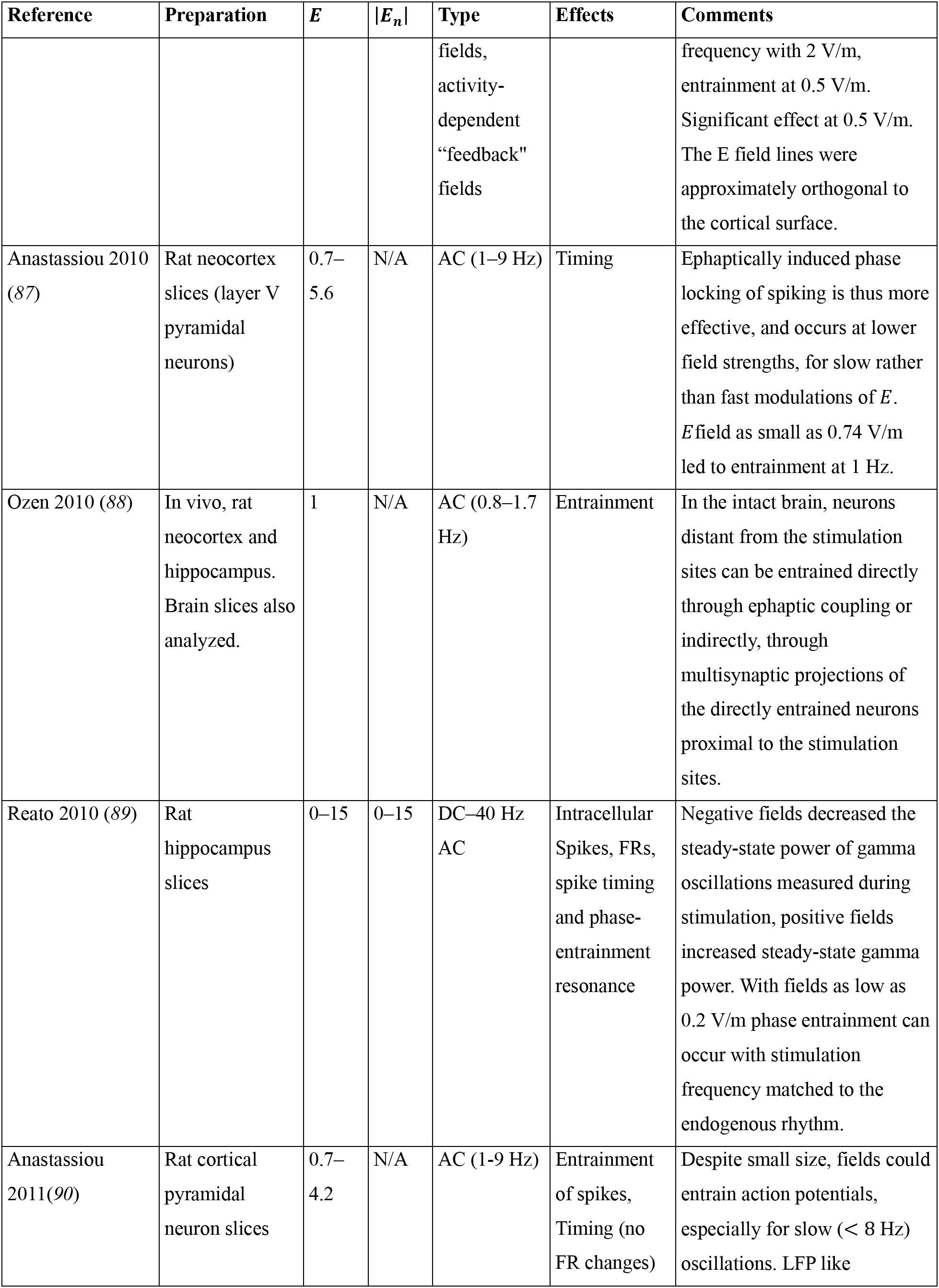
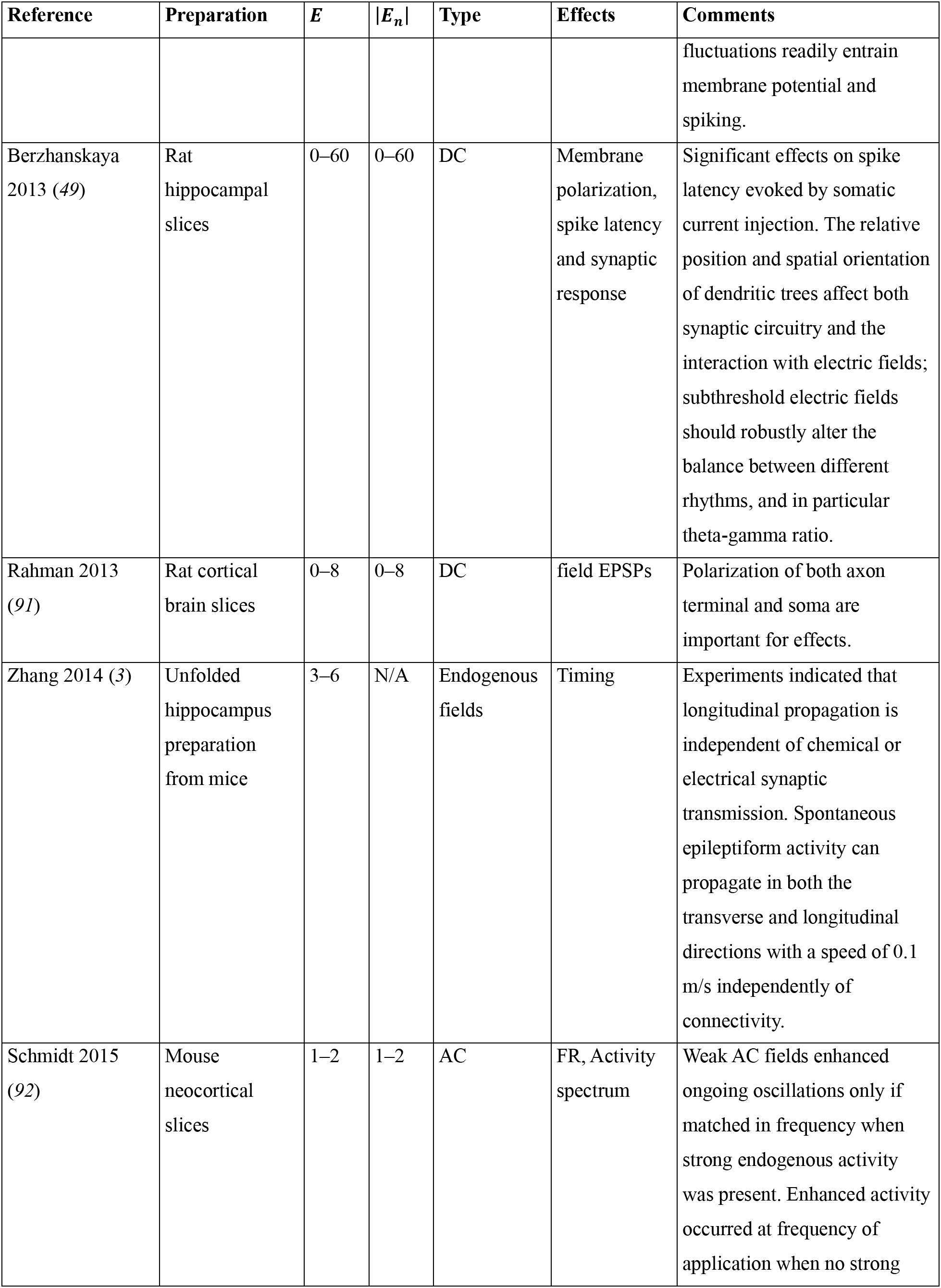
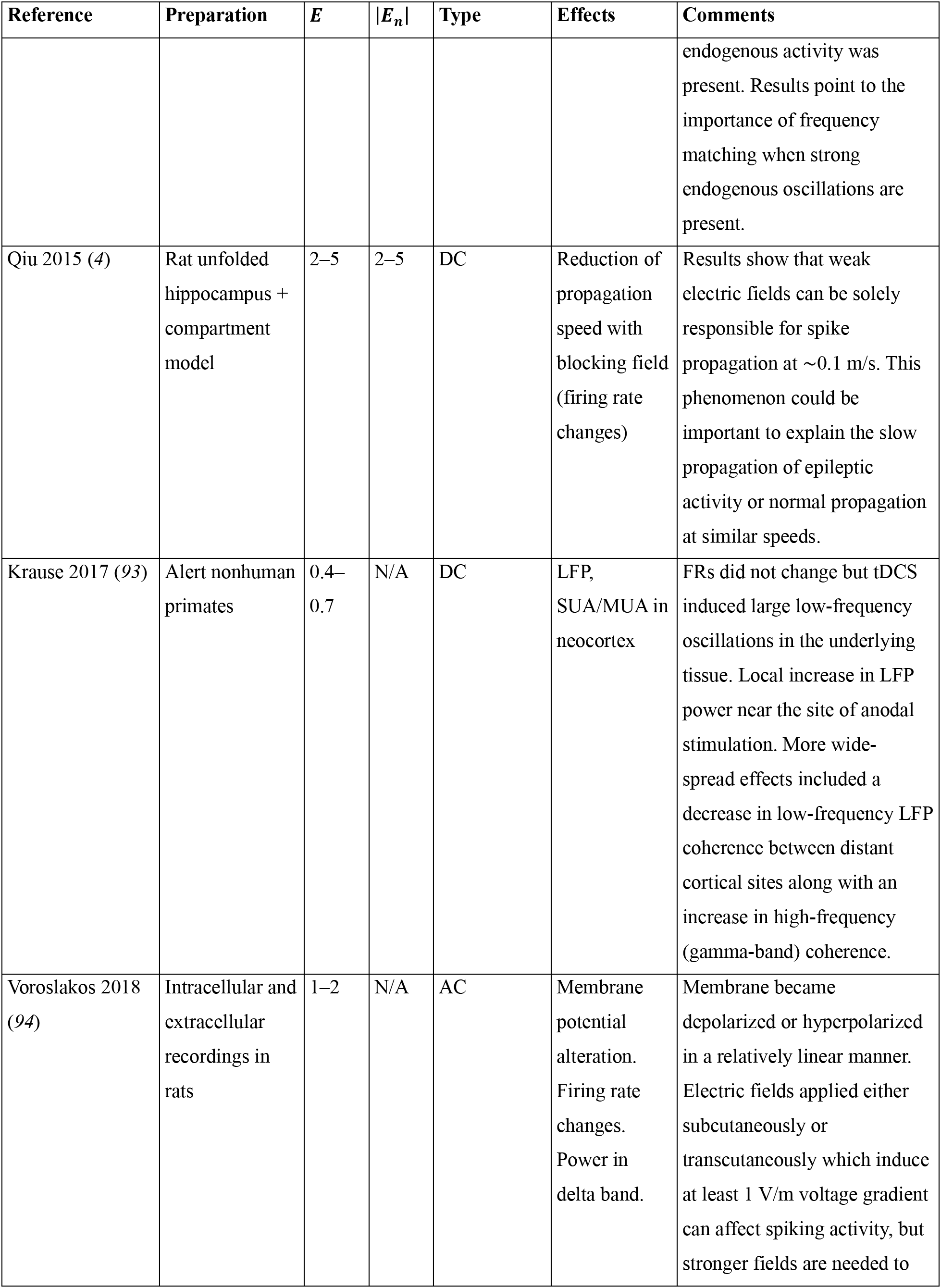
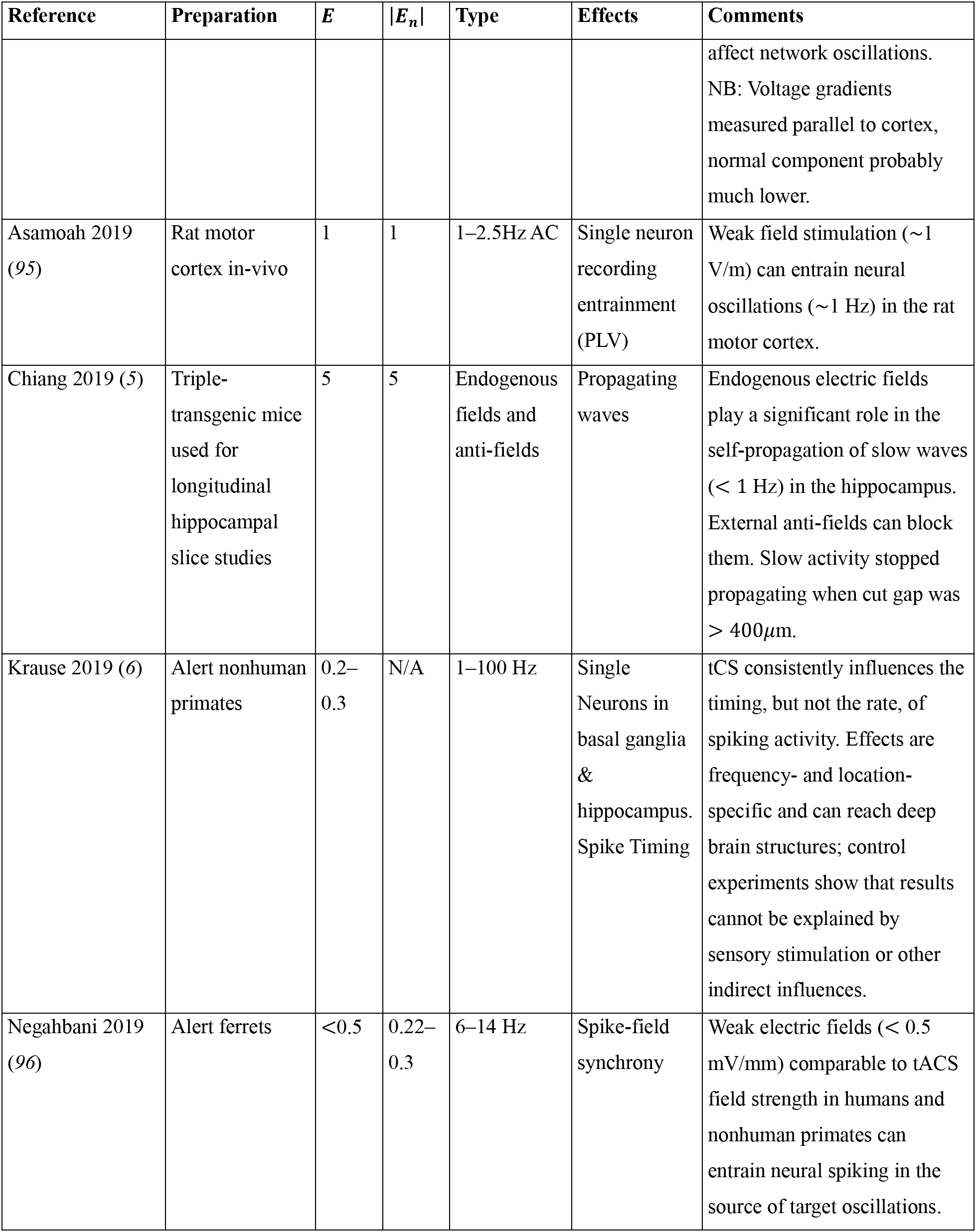
Overview of relevant work highlighting the physiological impact of weak electric fields in-vitro or in-vivo and providing quantitative measurements of electric field. The range of electric field magnitude (*E* = ||***E***||) or of the normal component of the electric field to cell layers (*E_n_*) in V/m (equivalently, mV/mm), that have been shown to influence function are listed. Only references where at least the magnitude of the extracellular electric field is specified are used (the voltage gradient). EPs: evoked potentials. AC: alternating current. DC: direct current. FR: firing rates. LFP: local field potential. SUA/MUA: single/multiple unit activity.

**Figure S1:**
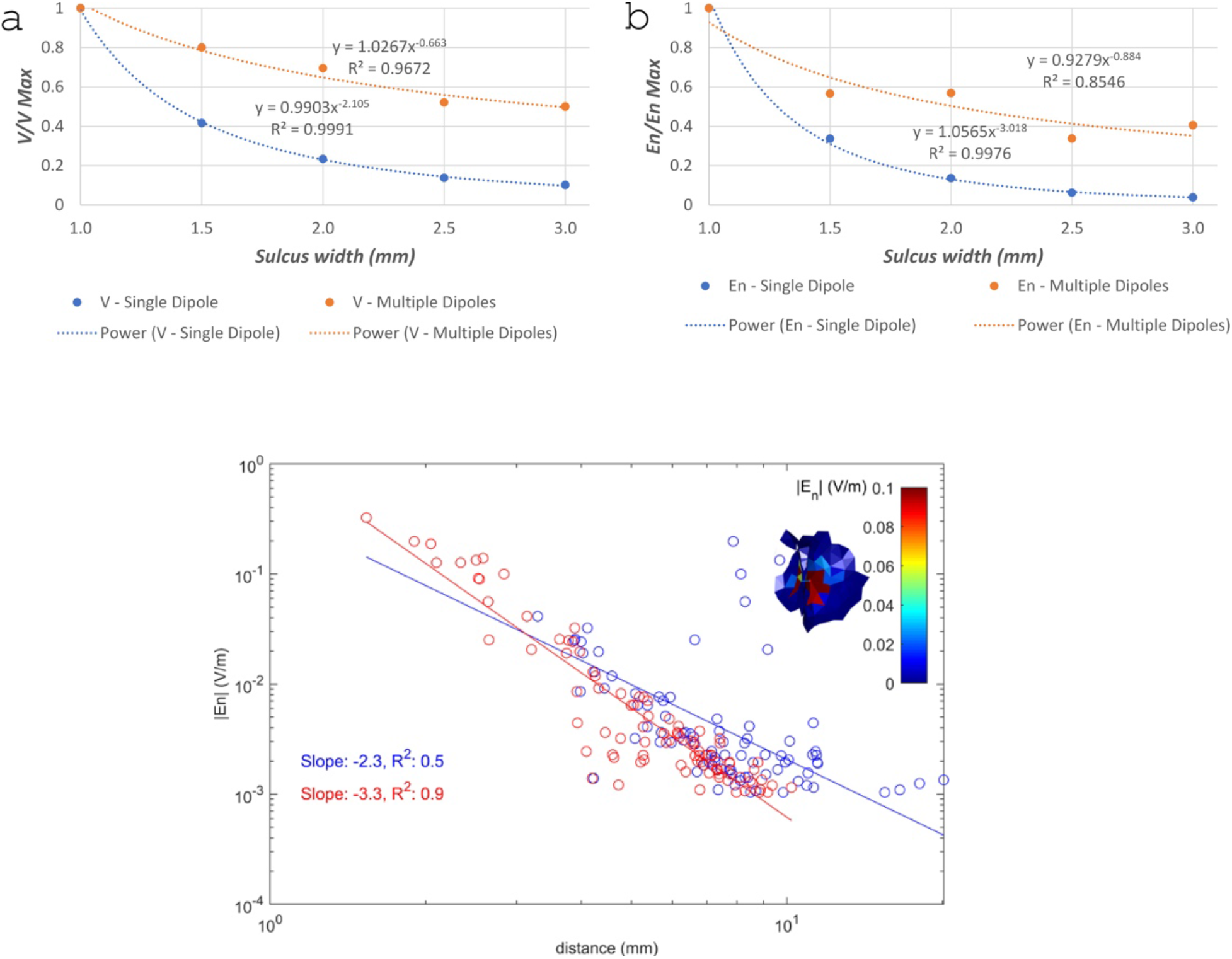
Decay of ***V*** and *E_n_* in the 2D and 3D models of the sulcus. Top: field decay in 2D model: (a) Decay of *V* with sulcus width in the single source model (blue dots) and multiple sources model (orange dots). The fit to a power function is also shown for each model. (b) Same as (a), but now for *E_n_*, the component of the electric field normal to the sulcus wall. Bottom: field decay in 3D model: l*oglog* plot of |*E_n_*| (in V/m) in the GM-CSF surface as a function of the logarithm of the geodesic (blue dots) or Euclidean (red dots) distance (in mm) to the dipole. The inset shows *E_n_* (in V/m) in a 3D rendering of the cortical surface. The location of the source is indicated by the red arrow. Only points where the absolute value of *E_n_* is between 0.001 V/m and 1.0 V/m are shown. Linear fits to these plots are also shown, together with the slope and *R*^2^ values.

**Figure S2:**
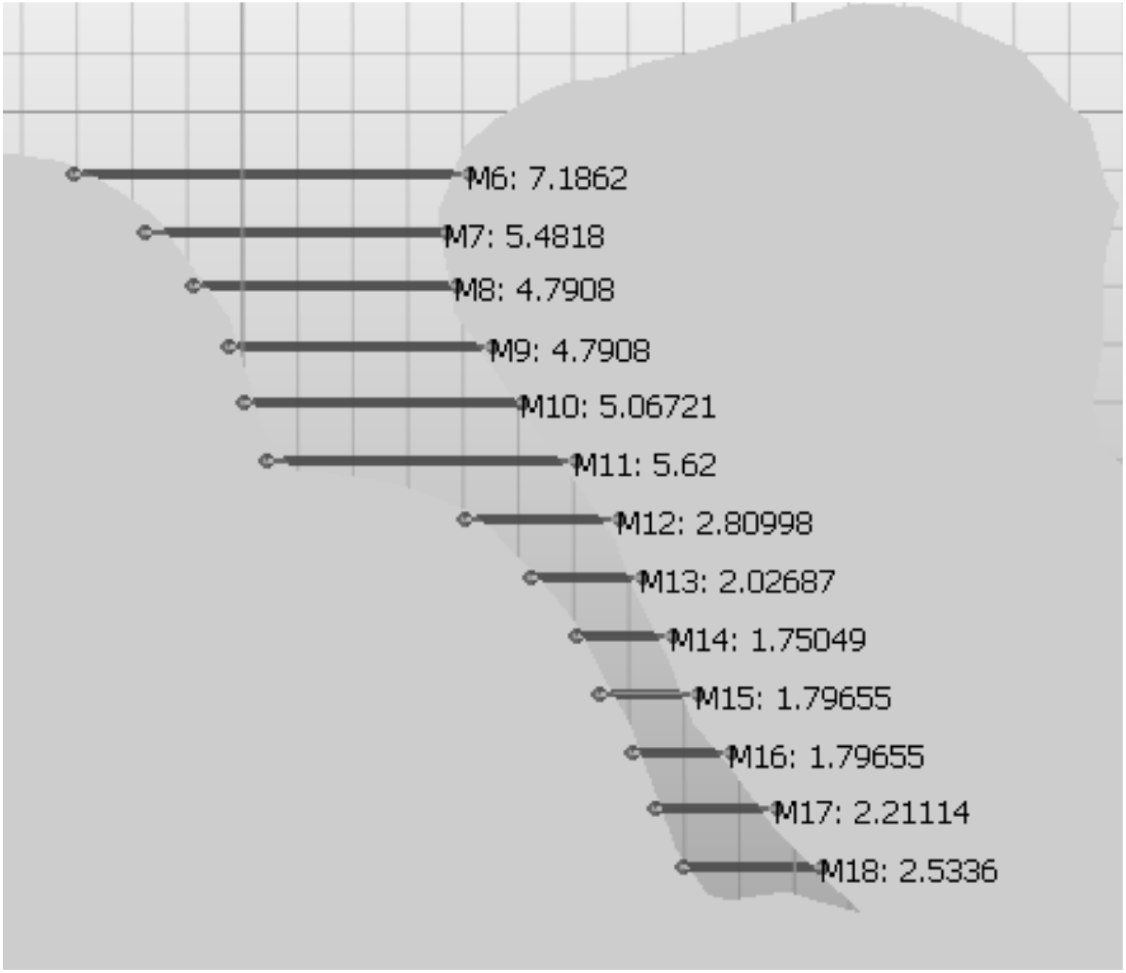
Sulcus geometry. Measurements of width (mm) in the sulcus used for realistic modeling in Figure 3 in the main text.

**Figure S3:**
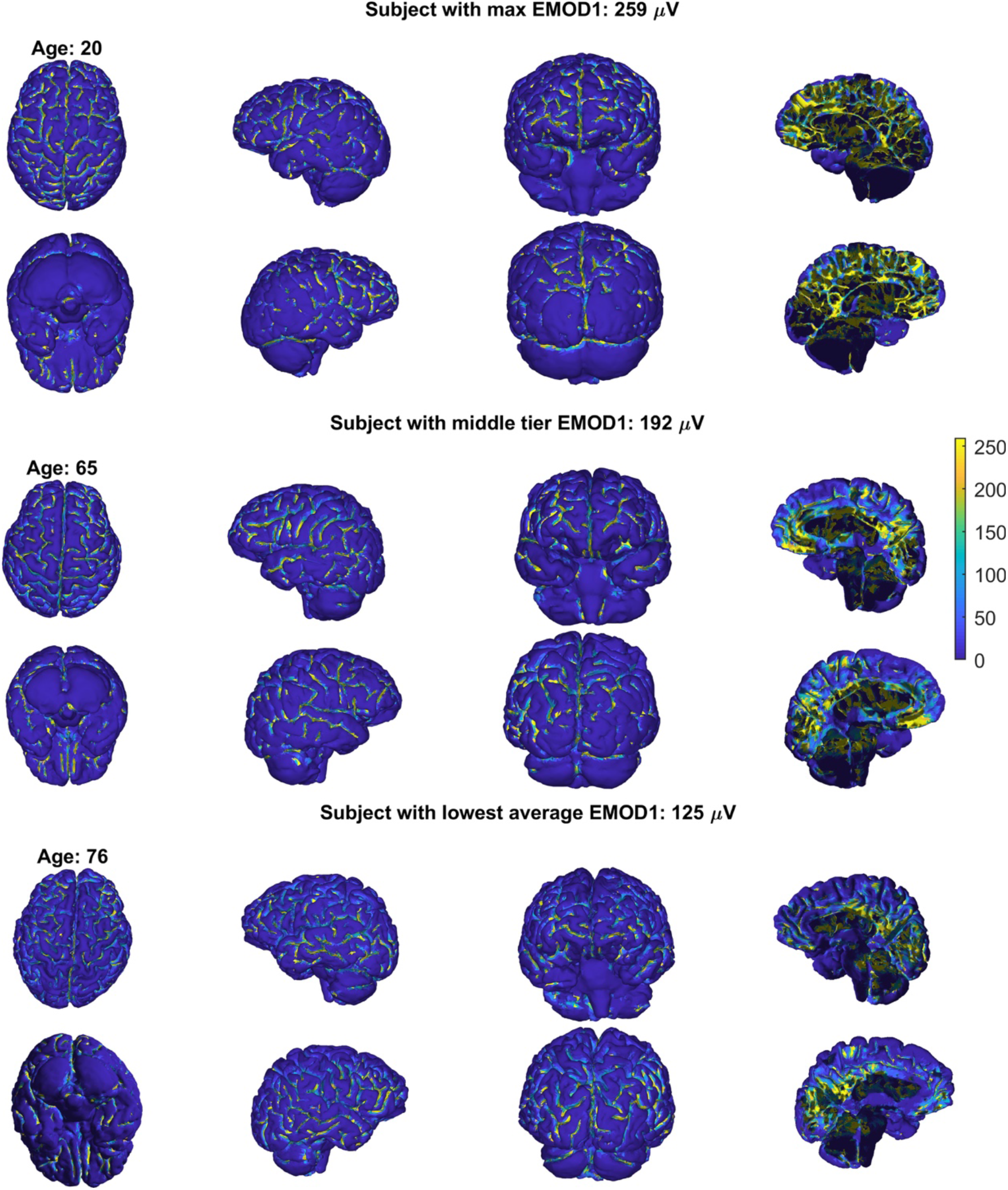
Surface distribution of the EMOD1 coefficient (*l*_0_ of 5 mm) for subjects with different ages. Subjects are presented from highest (top) to lowest EMOD1 (bottom) values. The color scale is common across all the plots. From left-right: top/bottom view, left/right-hemisphere view, front/back view, mid sagittal place left/right hemisphere view.

**Figure S4:**
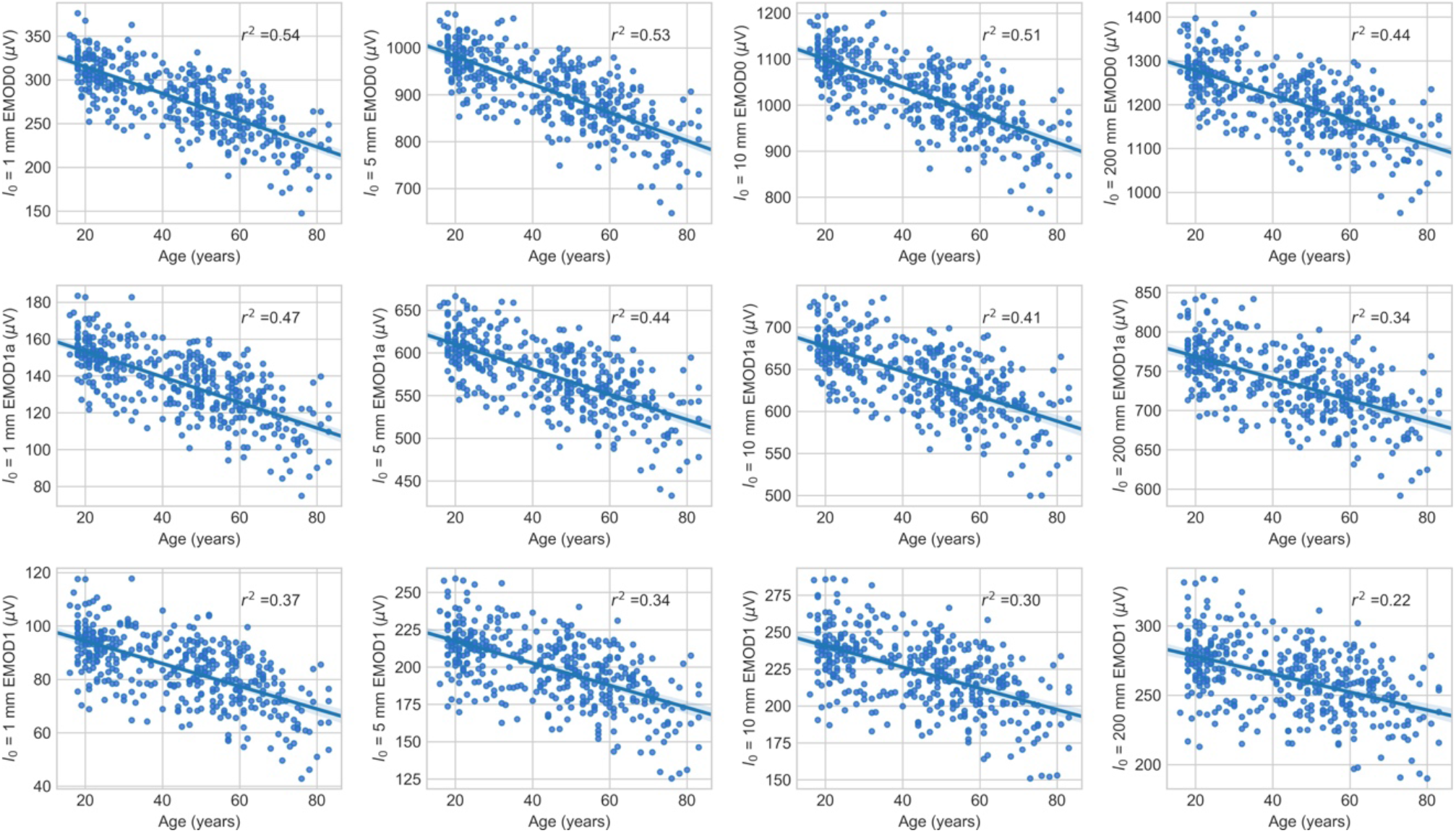
Linear fits of EMOD variants to age. Different rows correspond to different EMOD1 variants: EMOD0 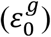, EMOD1a 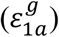 and EMOD1 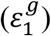. Different columns correspond to different 10 parameters: 1, 5, 10 and 200 mm, respectively from left to right.

**Figure S5:**
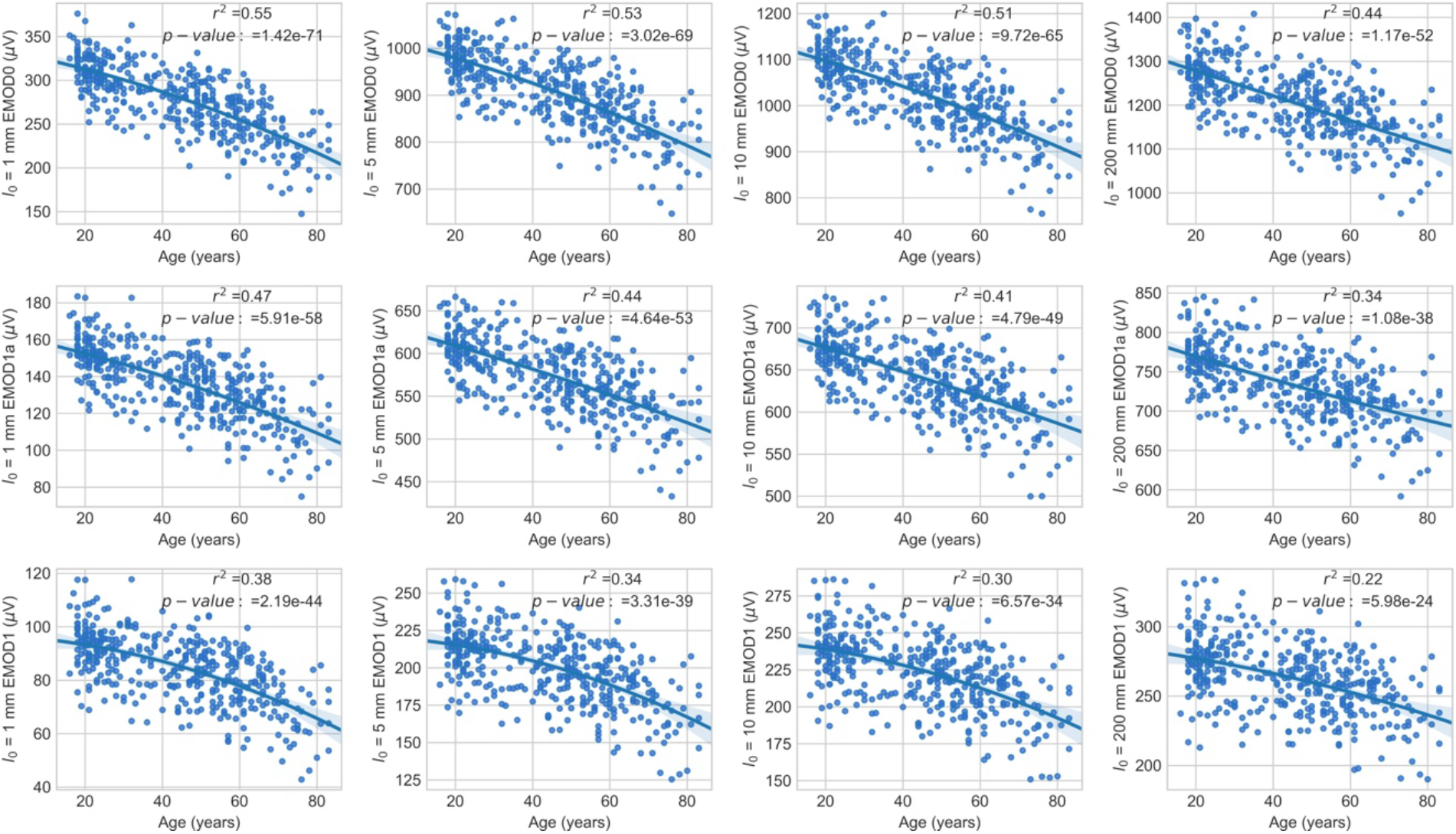
Second order fits of EMOD variants to age. Different rows correspond to different EMOD1 variants: EMOD0 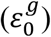, EMOD1a 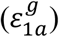 and EMOD1 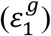. Different columns correspond to different *l*_0_ parameters: 1, 5, 10 and 200 mm, respectively from left to right.

**Figure S6:**
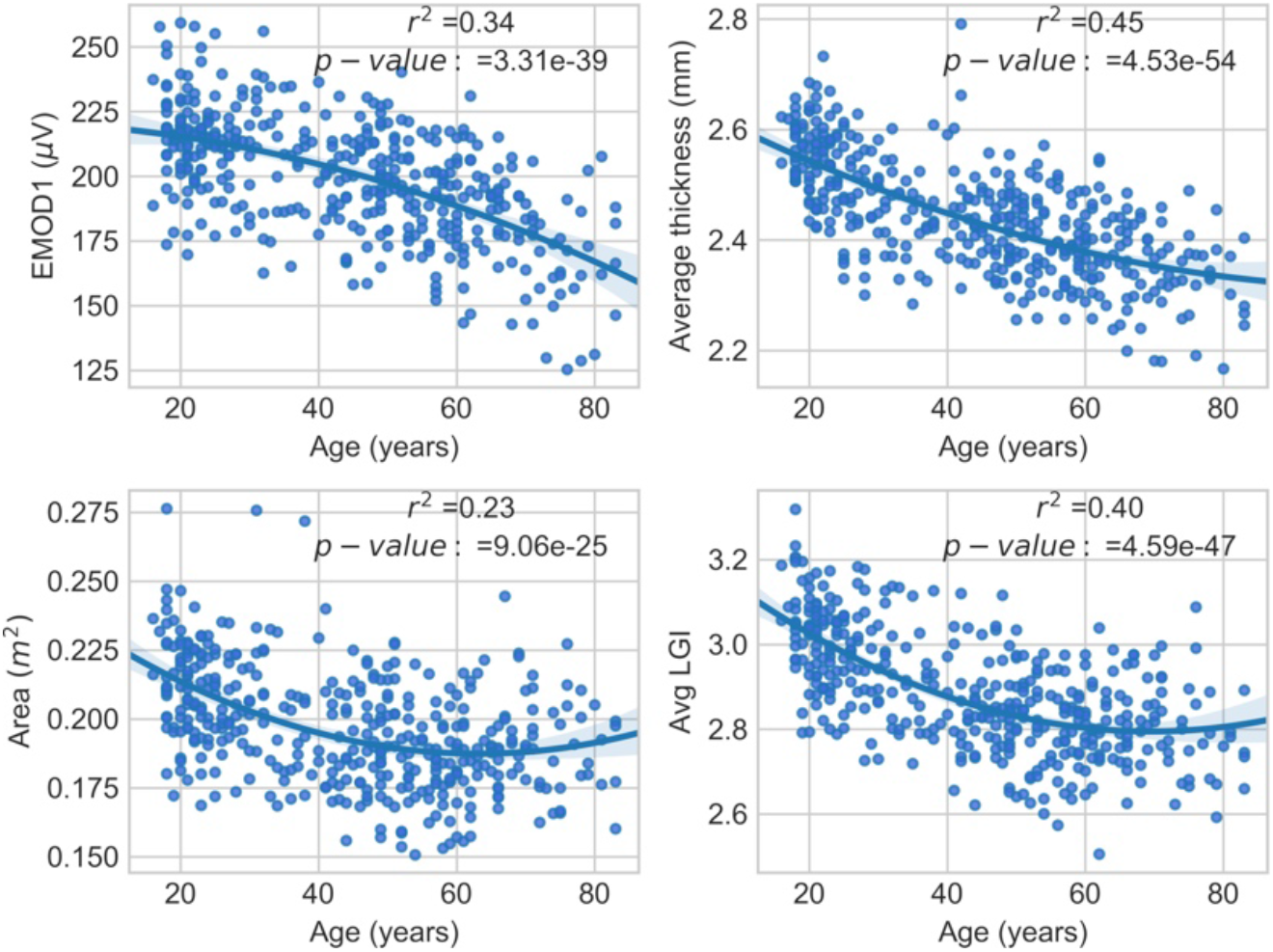
Second order fits of EMOD1, average LGI, average cortical thickness and cortical area to age. For each plot, r-squared and p-values for the fit are shown as well.

**Figure S7:**
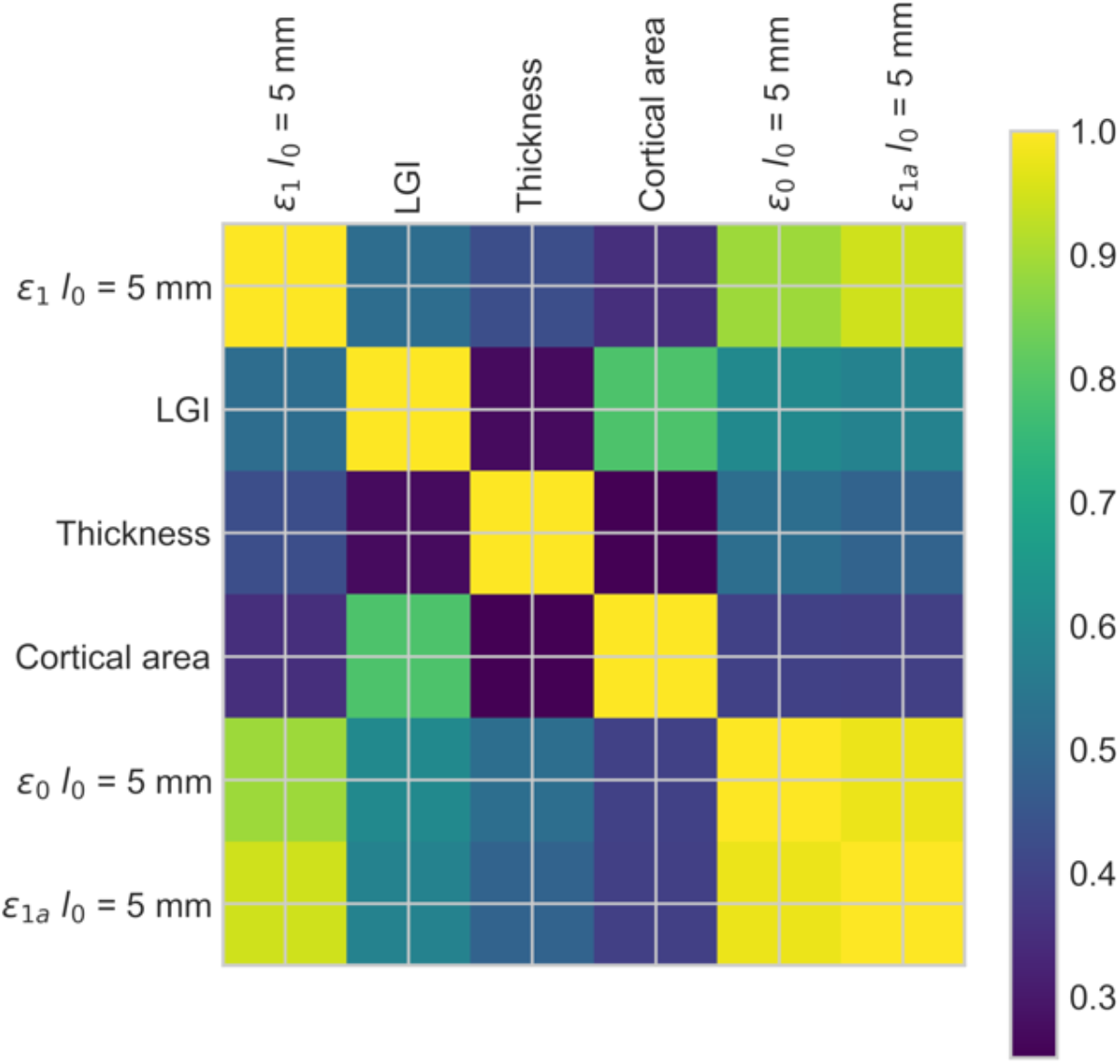
**Pearson correlation coefficients** between different EMOD variants, average LGI, average cortical thickness and total surface area.

**Figure S8:**
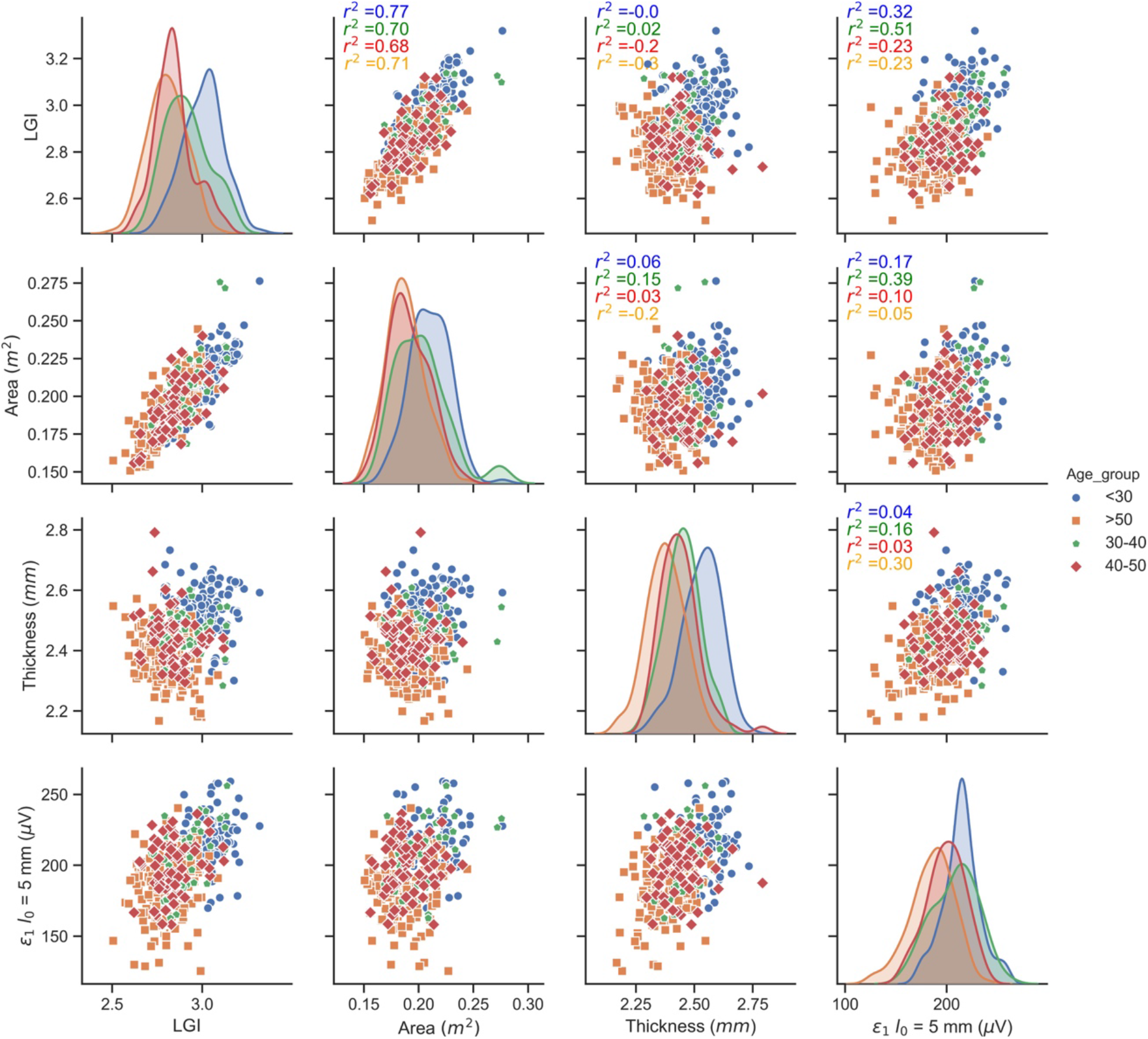
Correlation between average LGI, EMOD1 (*l*_0_ set to 5 mm), average cortical thickness and total cortical area for different age range groups. The plots along the main diagonal show histograms of these quantities grouped by age range. The offline range elements show each variable plotted against all others. Pearson correlation coefficients for each pairing, divided by age group, are also presented.

**Figure S9:**
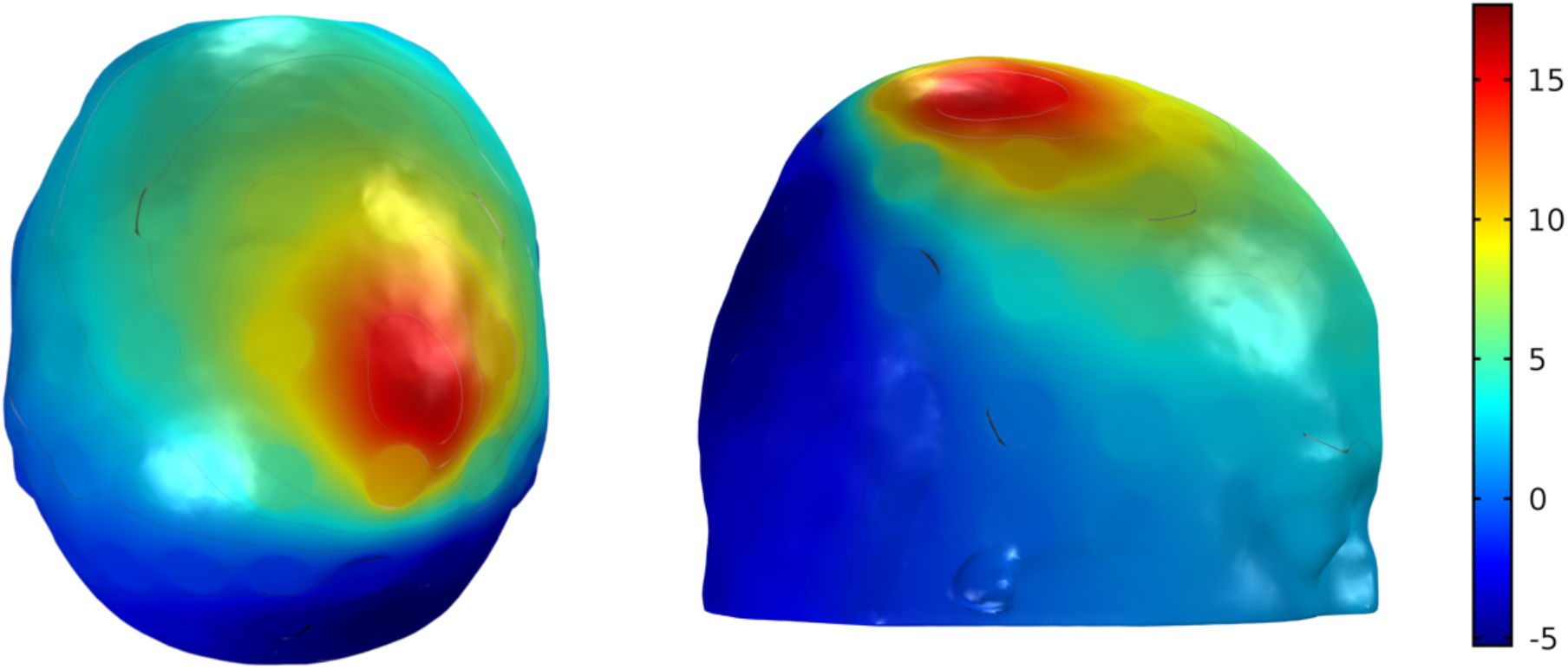
EEG (referenced to T8, in μV) as generated by cortical patch in Figure 3 (see also Table 1). The dipole patch consists of 133 dipole sources (patch area of 5.3 cm^2^), with a dipole density of 0.5 nAm/mm^2^.

